# Cell type-specific role of lamin-B1 and its inflammation-driven reduction in organ building and aging

**DOI:** 10.1101/448837

**Authors:** Sibiao Yue, Xiaobin Zheng, Yixian Zheng

**Author notes:** Co-first author Contact (Sibiao Yue, Xiaobin Zheng).

## Abstract

Cellular architectural proteins often participate in organ development and maintenance. Although functional decay of some of these proteins during aging is known, the cell-type specific developmental role and the cause and consequence of their subsequent decay remain to be established especially in mammals. By studying lamins, the nuclear structural proteins, we demonstrate that lamin-B1 functions specifically in the thymic epithelial cells (TECs) for proper thymus organogenesis. An upregulation of proinflammatory cytokines in the intra-thymic myeloid immune cells during aging accompanies a gradual reduction of adult TEC lamins-B1. These cytokines cause adult TEC senescence and lamin-B1 reduction. We identify 17 adult TEC subsets and show that TEC lamin-B1 maintains the composition of these TECs. Lamin-B1 supports the expression of TEC genes needed for maintaining adult thymic architecture and function. Thus, structural proteins involved in organ building and maintenance can undergo inflammation-driven decay which can in turn contribute to age-associated organ degeneration.

## Introduction

Cell and tissue architectural proteins, such as extracellular matrices, cytoskeleton, and nucleoskeleton, are involved in maintaining cell and tissue shapes. It is thus generally accepted that these proteins are required for tissue/organ building and their ‘wearing and tearing’ during the process of maintaining tissue/organ functions could contribute toward aging. However, due to their often ubiquitous presence in all cell types, not much effort has been devoted to understand whether they have cell-type specific roles in organ building, if organ aging is caused by the cell type specific decay of these proteins, and the cause of such decay during aging.

Among the structural proteins, lamins, the major component of the nuclear lamina that form a filamentous meshwork, have been implicated in proper organogenesis (Coffinier et al., 2011; Y. Kim et al., 2011). Interestingly, reduction of lamin-B1 is found in the aging human skins (Dreesen, Chojnowski, et al., 2013), Alzheimer’s Disease patient brains (Frost, Bardai, & Feany, 2016), and various *Drosophila* organs (H. Chen, Zheng, & Zheng, 2014; Tran, Chen, Zheng, & Zheng, 2016), but the cause of such reduction and its impact on organ function, especially in mammals, remains poorly understood.

Elevated proinflammatory cytokines in aging animals, including humans, have been shown to contribute to various organ dysfunctions and human diseases (Franceschi et al., 2000). Indeed, extensive studies *in vitro* have shown that proinflammatory cytokines can induce senescence of a number of tissue culture cells (Acosta et al., 2008; Dumont, Balbeur, Remacle, & Toussaint, 2000; Kuilman et al., 2008). For example, either overexpression of CXCR2 in human primary fibroblasts or treatment of these cells with IL-1α or TNF-α induces cellular senescence (Acosta et al., 2008; Dumont et al., 2000). These proinflammatory cytokines can also reinforce cellular senescence in other primary tissue culture cells triggered by forced oncogene expression (Kuilman et al., 2008). Despite these studies, however, the cell/tissue source of age-associated inflammation and whether such inflammation disrupts structural proteins and thus contributes to organ aging remain unclear in any organism.

Considering the varied environments different tissues/organs reside in and the different functions they perform, it is highly likely that the inflammatory causes and consequences are different in different tissues and organisms. Cellular senescence triggered by inflammation has been implicated in aging and organ degeneration in mammal (Ren et al., 2009). The multitudes of senescence-associated cellular changes have, however, made it difficult to pinpoint which of these changes makes a key contribution toward age-associated organ dysfunction. Additionally, vertebrate organs often contain complex cell types, which makes it challenging to identify the cell source(s) and target(s) of inflammation that contribute to organ aging. Among many organs, the vertebrate thymus has a relatively simple stromal cell population called thymic epithelial cells (TECs) that are essential for thymic development, organization, and function (G. Anderson & Takahama, 2012). The TECs can thus serve as a relatively simple model to understand how inflammation and cellular senescence could influence structural proteins and in turn contribute to organ aging.

As a primary lymphoid organ, the thymus produces naïve T cells essential for adaptive immunity. Differentiated from the Foxn1 positive progenitors, the TECs consist of cortical TECs (cTECs) and medullary TECs (mTECs) that make up the cortical and medullary compartments of the thymus, respectively (Boehm, Nehls, & Kyewski, 1995). Whereas the cTECs play a major role in the positive selection of T cells, the mTECs along with the thymic dendritic cells (DCs) mediate central tolerance by facilitating clonal deletion of self-reactive T cells (G. Anderson & Takahama, 2012; Corbeaux et al., 2010).

The age-associated thymic involution or size reduction is known to contribute to the dysfunction of the immune system (Chinn et al., 2012). Studies in mice have shown that thymic involution can be separated into two phases (Aw et al., 2008; Aw & Palmer, 2012; Shanley et al., 2009). The first phase occurs within ~6 weeks after birth and is characterized by a rapid reduction of thymic size. This phase is referred to as the developmentally related involution and it does not negatively affect the immune system. The second phase of thymic involution occurs during the process of organism aging and is manifested as a gradual reduction of thymic size and naïve T cell production. Foxn1 reduction in TECs soon after birth appears to contribute to the first developmental phase of thymic involution (L. Chen, Xiao, & Manley, 2009; O’Neill et al., 2016; Rode et al., 2015), but the cause of the second and age-associated phase of involution is unknown.

We show that of the three lamins, only lamin-B1 is required in TECs for the development and maintenance of the spatially segregated cortical and medulla compartments critical for proper thymic function. We identify several proinflammatory cytokines in aging thymus that trigger TEC senescence and TEC lamin-B1 reduction. Importantly, we report the identification of 17 adult TEC subsets and show that lamin-B1 reduction in postnatal TECs contributes to the age-associated TEC composition change, thymic involution, reduced naïve T cell production, and lymphopenia.

## Results

### Lamin-B1 functions in TECs to support proper thymic development

To understand if lamins play a role in thymus development, we first analyzed the thymuses in mice deleted of the lamin-B1 gene, *Lmnb1*, in the germline and their littermate controls during embryogenesis. We found that the embryonic (E) day 18.5 (E18.5) *Lmnb1* null thymuses were smaller than the age-matched littermate controls (Figure 1A). Since defects in thymocytes or TECs or both can result in small thymuses, we sought to identify the cell type where *Lmnb1* plays an important role during thymic organogenesis. We used the *Lmnb1^f/f^* allele derived from *Lmnb1*^tm1a(EUCOMM)Wtsi^ from the EUCOMM project. These mice were crossed with the *Lck-Cre* or *Foxn1-Cre* (*FN1Cre*) mice to generate T-cell- or TEC-specific *Lmnb1* knockout mice, respectively (Gordon et al., 2007; Hennet et al., 1995). We found that ablation of *Lmnb1* in TECs resulted in a significant reduction in the size and total cell number in the 2-month (mon)-old mouse thymuses compared to the littermate controls, whereas deleting *Lmnb1* in T cells had no apparent effect (Figure 1B and C). Since *FN1Cre* begins to express Cre at E11.5 in TECs (Gordon et al., 2007), we analyzed thymuses in mice from E18.5 to 2 mon after birth. We found that the thymuses isolated from the *Lmnb1^f/f^;FN1Cre* mice were smaller in size and had reduced total cell numbers as compared to the littermate controls, whereas *Lmnb1* deletion in the *Lmnb1^f/f^;LckCre* mice had no such effect (Figure 1C).

**Figure 1.**
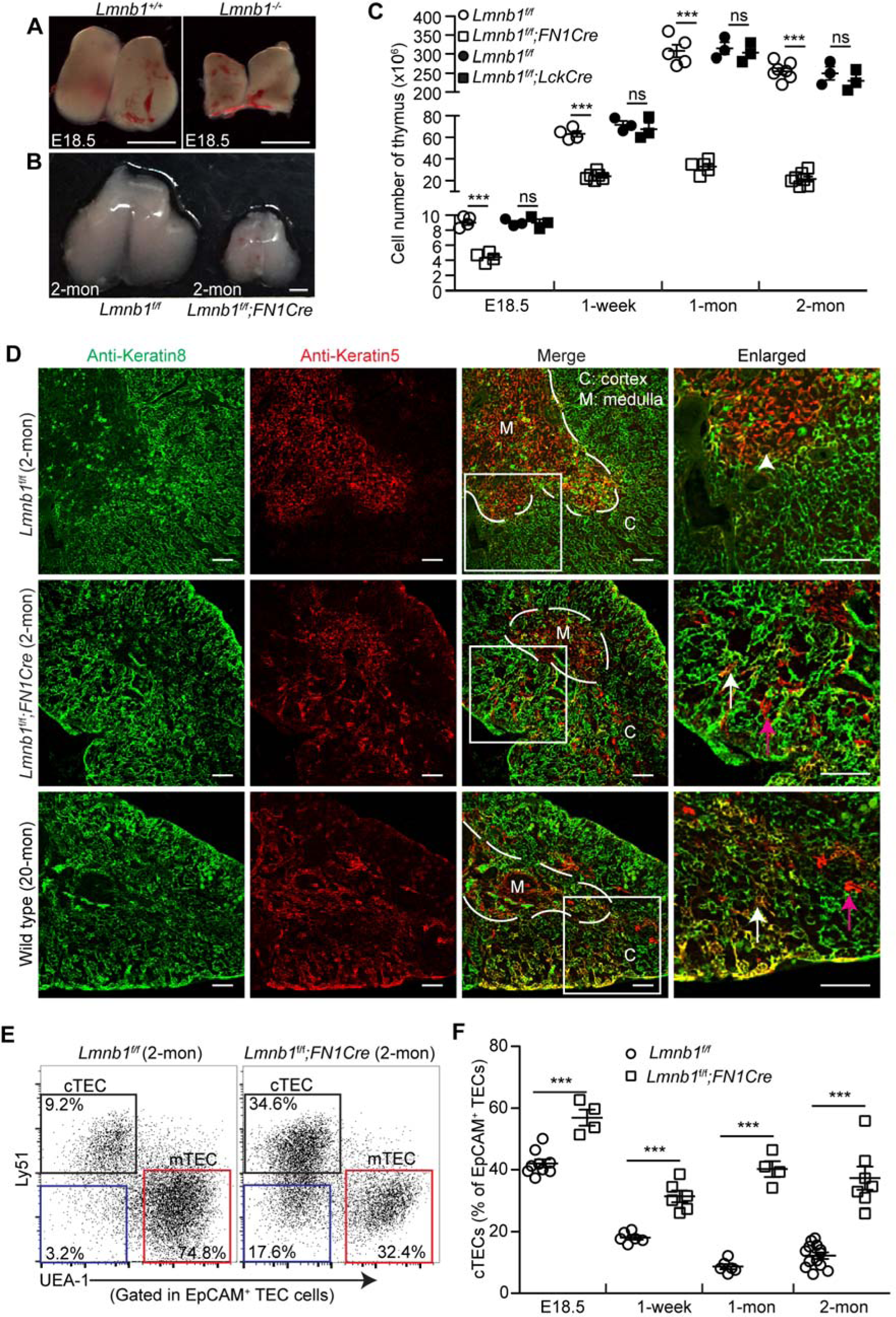
Effects of embryonic *Lmnb1* deletion in TECs on thymic organogenesis. (A-B) Representative images of thymuses from embryonic day 18.5 (E18.5) *Lmnb1*^*+/+*^ and *Lmnb1*^*-/-*^ mice (A) or 2-mon-old *Lmnb1*^*f/f*^ and *Lmnb1^f/f^;Foxn1Cre* (*FN1Cre*) mice (B). Scale bars, 1mm. (C) Total thymic cell number counts from mice with the indicated ages and genotypes. (D) Distribution of cTECs (K8, green) and mTECs (K5, red) in thymuses with indicated genotypes as revealed by immunostaining. A 20-mon-old WT thymus is shown as an aged thymus control. White dash lines demarcate the cortical and medulla junction regions (CMJ) in the thymuses. A section of thymuses (white squares) in each genotype is enlarged to the right with the white arrowhead marking the K5^+^K8^+^ TECs in the CMJ in the 2-mon-old *Lmn1*^*f/f*^ control thymus and the arrows marking the spreading of K5^+^K8^+^ TECs (white) and K5^+^ TECs (purple) into the cortical regions in the other thymuses. Scale bars, 100 >m. (E) Representative flow cytometric profiles showing the frequency of cTECs (Ly51^+^UEA-1^-^, black squares) and mTECs (UEA-1^+^Ly51^-^, red squares) from 2-mon-old *Lmnb1*^*f/f*^ and *Lmnb1^f/f^;FN1Cre* thymuses. The displayed cells (each dot represents one cell) are gated first on CD45^-^EpCAM^+^ cells and then analyzed according to Ly51 and UEA-1 to identify cTEC and mTEC subsets. The percentage of UEA-1^-^ Ly51^-^ TECs is shown in the bottom left corner (blue squares). (F) Summary of the frequency of cTECs in *Lmnb1*^*f/f*^ control and *Lmnb1^f/f^; FN1Cre* thymuses at the indicated ages. Each circle or square represent one control *Lmnb1*^*f/f*^ and one *Lmnb1^f/f^;FN1Cre* mouse, respectively. Error bars, standard error of the mean (SEM) based on at least three independently analyzed mice. Student’s t test: *P<0.05, ** P<0.01, ***P<0.001; ns, not significant. See also Figure S1.

We next assessed the role of the other two lamins, lamin-B2 encoded by *Lmnb2* and lamin-A/C encoded by *Lmna,* in TECs. The conditional *Lmnb2* allele, *Lmnb2^f/f^*, was derived from *Lmnb2*^tm1a(KOMP)Wtsi^ (KOMP project) by breeding with ACTB-FLPe mice, which express FLP1 recombinase in a wide variety of cells including germ cells, to remove the neomycin cassette flanked by Frt sites. The *Lmna^f/f^* we generated in house was reported previously (Y. Kim & Zheng, 2013). The *Lmnb2^f/f^* or *Lmna^f/f^* mice were crossed with *FN1Cre* mice. Genotyping and qPCR analyses revealed a similar deletion efficiency of all three lamin genes by *FN1Cre* in both cTECs and mTECs (Figure S1A-C). We found that depletion of either *Lmnb2* or *Lmna* in TECs had no obvious effect on thymic size and total cellularity (Figure S1D-F). These analyses show that lamin-B1, but not lamin-B2 and lamin-A/C, in TECs supports the proper building of thymus during development.

### Lamin-B1 supports proper development of cortical and medulla thymic compartments

To analyze whether lamin-B1 deletion affects the organization of TECs, we immunostained cTECs and mTECs using anti-keratin 8 (K8) or 5 (K5) antibodies, respectively, in the thymuses from the 2-mon-old *Lmnb1^f/f^;FN1Cre* mice or their littermate controls. We found that lamin-B1 deletion in TECs resulted in an intermingling of the cortical and medullary TEC compartments compared to the controls (Figure 1D). Interestingly, this intermingling of the two TEC compartments are highly reminiscent of the mixed cTEC and mTEC compartments known to occur in the old WT mouse thymuses (Figure 1D). The breakdown of the cortical and medullary compartments is reflected in an increased presence of K5 and K8 double-positive (K5^+^K8^+^) TECs throughout thymuses and the presence of K5^+^ mTECs outside of the medullary region of both the 2-mon-old *Lmnb1^f/f^;FN1Cre* and the aged 20-mon-old WT mouse thymuses, whereas in the young WT thymuses the K5^+^K8^+^ TECs were located in the cortical and medullary junction (CMJ) regions (Figure 1D).

The EpCAM^+^TECs can be further separated into mTECs and cTECs as each subset of cells are either UEA-1^+^ or Ly51^+^, respectively. Using flow cytometry, we further analyzed the mTECs (EpCAM^+^UEA-1^+^Ly51^-^) and cTECs (EpCAM^+^UEA-1^-^Ly51^+^) and found a pronounced reduction in the frequency of mTECs and a skewing of the TEC population toward cTECs in the *Lmnb1^f/f^;FN1Cre* thymuses compared to the littermate controls (Figure 1E and F). There was also a marked increase of a TEC subpopulation that did not express the canonical surface UEA-1 or Ly51 (Figure 1E, blue squares) in the *Lmnb1^f/f^;FN1Cre* thymuses. These analyses demonstrate that lamin-B1 is not required for the TEC lineage commitment but it plays a role in efficient TEC differentiation and TEC compartment organization during thymus organogenesis.

### Embryonic lamin-B1 deficiency in TECs impairs positive selection of conventional αβT-cells

Since the disrupted mTEC and cTEC compartmentalization and composition upon laminB1 deletion could affect proper TEC niche formation, thereby affecting T-cell development (thymopoiesis) (G. Anderson & Takahama, 2012), we examined the impact of lamin-B1 deficiency in TECs on thymopoiesis. We first analyzed the CD4 and CD8 single positive (SP) thymocytes and found a 50-60% reduction in each of these thymocytes in the 2-mon-old *Lmnb1^f/f^;FN1Cre* thymuses compared to the littermate controls (Figure 2A and B). Flow cytometry analyses of cell surface expression of TCRβ chain and CD69 on thymocytes further revealed an ~50% reduction of the TCRβ^high^CD69^+^ and mature TCRβ^high^CD69^-^populations in the 2-mon-old *Lmnb1^f/f^;FN1Cre* (Figure 2C and D). We further confirmed this finding by using the OT-II transgenic TCR mouse model and found that depletion of *Lmnb1* by *FN1Cre* led to a substantial reduction of the transgenic OT-II CD4^+^SP thymocytes (Figure 2E and F).

**Figure 2.**
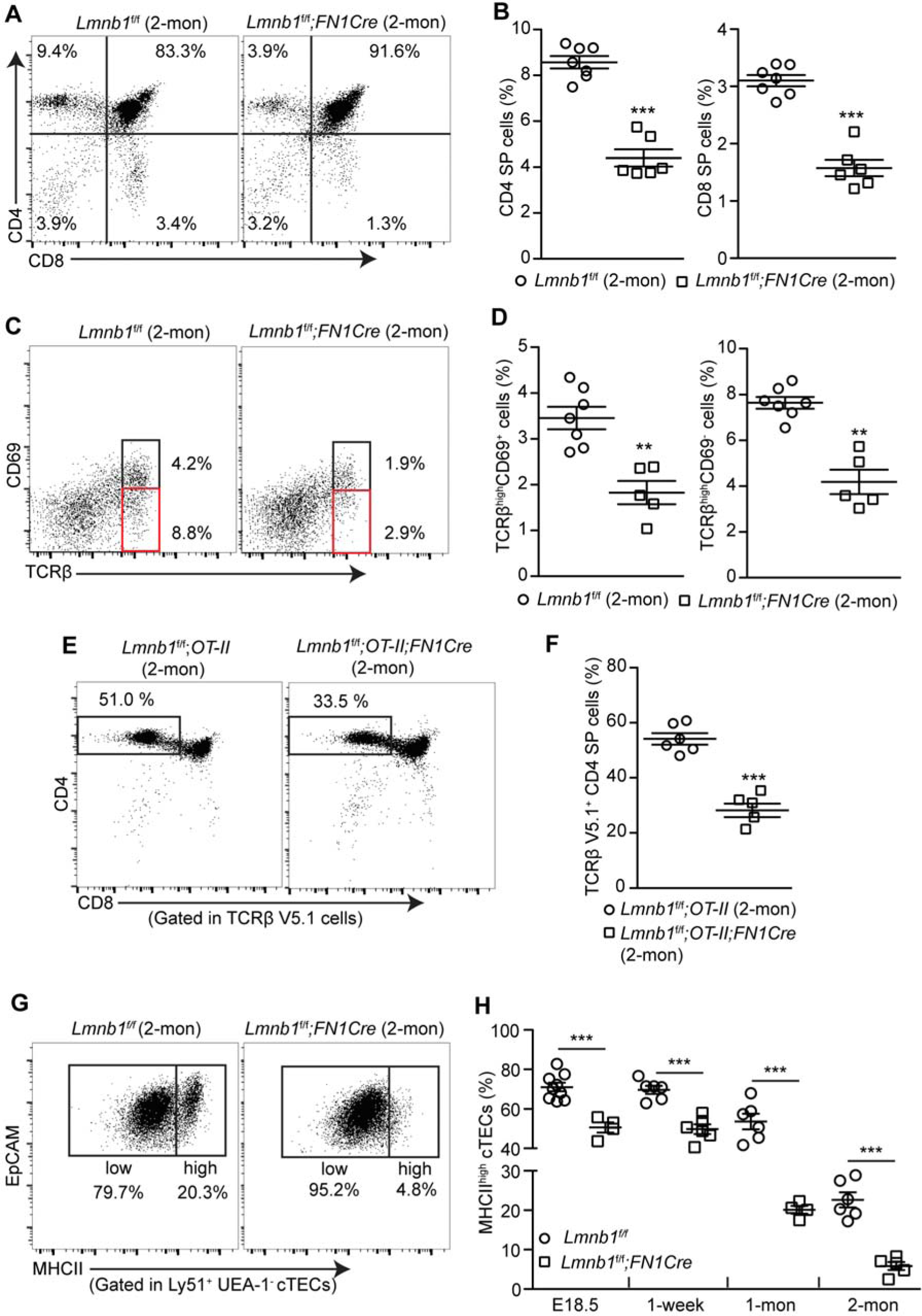
Effects of developmental TEC *Lmnb1* deletion on the positive selection of αβT-cells in mouse thymus. (A) Representative flow cytometry profiles showing a reduction of CD4^+^ single positive (CD4^+^ SP) and CD8^+^ SP thymocytes in the 2-mon-old *Lmnb1^f/f^;FN1Cre* mouse thymuses compared to that of the *Lmnb1*^*f/f*^ controls. (B) Quantifications of the frequency of CD4^+^ and CD8^+^ SP thymocytes within the gated live cells in the 2-mon-old *Lmnb1*^*f/f*^ control (n=7) and *Lmnb1^f/f^;FN1Cre* (n=6) mouse thymuses. (C) Flow cytometry analysis of TCRβ and CD69 expression on thymocyte subsets undergoing positive selection in 2-mon-old *Lmnb1*^*f/f*^ control and *Lmnb1^f/f^;FN1Cre* mouse thymuses. Black and red boxes show the TCRβ^high^CD69^+^ or TCRβ^high^CD69^-^ thymocytes. (D) Quantification of the frequency of the TCRβ^high^CD69^+^ and TCRβ^high^CD69^-^cells in the 2-mon-old *Lmnb1*^*f/f*^ control (n=7) and *Lmnb1^f/f^;FN1Cre* (n=5) mouse thymuses. (E) Flow cytometry analysis showing a reduction of CD4^+^ SP thymocytes in the 2-mon-old *Lmnb1^f/f^;OT-II;F1NCre* thymuses (n=5) compared to the *Lmnb1^f/f^;OT-II* thymuses (n=6). (F) Quantification of frequency of CD4^+^ SP thymocytes within the gated OT-II transgenic TCR (TCRβ V5.1) thymocytes in experiments shown in (E). (G) Representative flow cytometry plots of staining for MHCII^high^ and MHCII^low^ cTECs from 2-mon-old *Lmnb1*^*f/f*^ and *Lmnb1^f/f^;FN1Cre* thymuses. (H) A summary showing the reduction of the most differentiated MHCII^high^ cTECs in the *Lmnb1^f/f^;FN1Cre* thymuses compared to the *Lmnb1*^*f/f*^ control thymuses at the indicated ages. Each circle or square represents one *Lmnb1*^*f/f*^ control or *Lmnb1^f/f^;FN1Cre* mouse, respectively. Error bars, SEM from at least four independently analyzed mice. Student’s t test: *P<0.05, **P<0.01, ***P <0.001; ns, not significant. See also Figure S2.

Next, we further studied how lamin-B1 deficiency in TECs can affect αβT-cell development. TECs can be separated into the MHCII^low^ immature and the MHCII^high^ mature and more differentiated TECs. A decline in the number of mature MHCII^high^TECs is a hallmark of the age-associated thymus change in mice (Chinn et al., 2012). Strikingly, we found an ~5-fold reduction of the mature MHCII^high^ cTECs, in the 2-mon-old *Lmnb1^f/f^;FN1Cre* mice compared to littermate controls (Figure 2G and H). A reduction, albeit less pronounced, in the frequency of the MHCII^high^ mature mTECs was also observed in the 2-mon-old *Lmnb1^f/f^;FN1Cre* mice (*Lmnb1^f/f^*: 54 ± 4%; *Lmnb1^f/f^;FN1Cre:*42 ± 3%; P value=0.003). Thus lamin-B1 plays a critical role in TECs, particularly in cTECs, to promote efficient TEC differentiation and maturation from their progenitors, thereby supporting the organization of thymic cortical and medulla compartments and αβT-cell generation.

Interestingly, the frequencies of the double-negative (DN) thymocytes and the γδT cells, whose development are not dependent on the expression of MHCII in cTECs (Chien et al., 2014), are similar in the 2-mon-old *Lmnb1^f/f^;FN1Cre* and the littermate control thymuses (Figure S2). These results suggest that lamin-B1 is dispensable in cTECs to support early stages of T-cell development.

### Gradual lamin-B1 reduction in TECs during age-associated thymic involution

Since the disruption of TEC compartments and T cell selection in the 2-mon-old *Lmnb1^f/f^;FN1Cre* thymuses resembled the changes in the old mouse thymus, we asked whether lamin-B1 reduction may accompany the age-associated thymic involution. The mouse thymus first undergoes a rapid phase of involution between six to eight weeks after birth followed by a second age-associated phase of gradual involution. We immunostained lamin-B1, -B2, and -A/C, in the CD4^+^CD8^+^double-positive (DP) thymocytes, cTECs and mTECs isolated from 2- and 20-mon-old WT mouse thymuses and analyzed the protein levels using flow cytometry. Quantification of the mean fluorescence intensity (MFI) revealed that lamin-B1 was reduced by >65% in cTECs and mTECs, but not in DP thymocytes, in old WT mice (Figure 3A and B). The lamin-B2 and lamin-A/C levels remained the same upon aging in these cells (Figure 3C). To further confirm this, we used fluorescence-activated cell sorting (FACS) to isolate cTECs and mTECs. We mixed an adherent keratin-negative cell line (RAW264.7 from ATCC #TIB-71) with the sorted cTECs or mTECs to control for variability of immunostaining (Figure 3D and E). By quantifying lamin-B1 intensity in individual TECs and normalizing against the lamin-B1 level of the spike-in RAW264.7 cells, we found that ~80% cTECs or mTECs exhibited >50% lamin-B1 reduction in the old thymus (Figure 3F). Western blotting analyses of the isolated TECs also showed a large reduction of lamin-B1 level upon aging, whereas lamin-A/C remained unchanged (Figure 3G).

**Figure 3.**
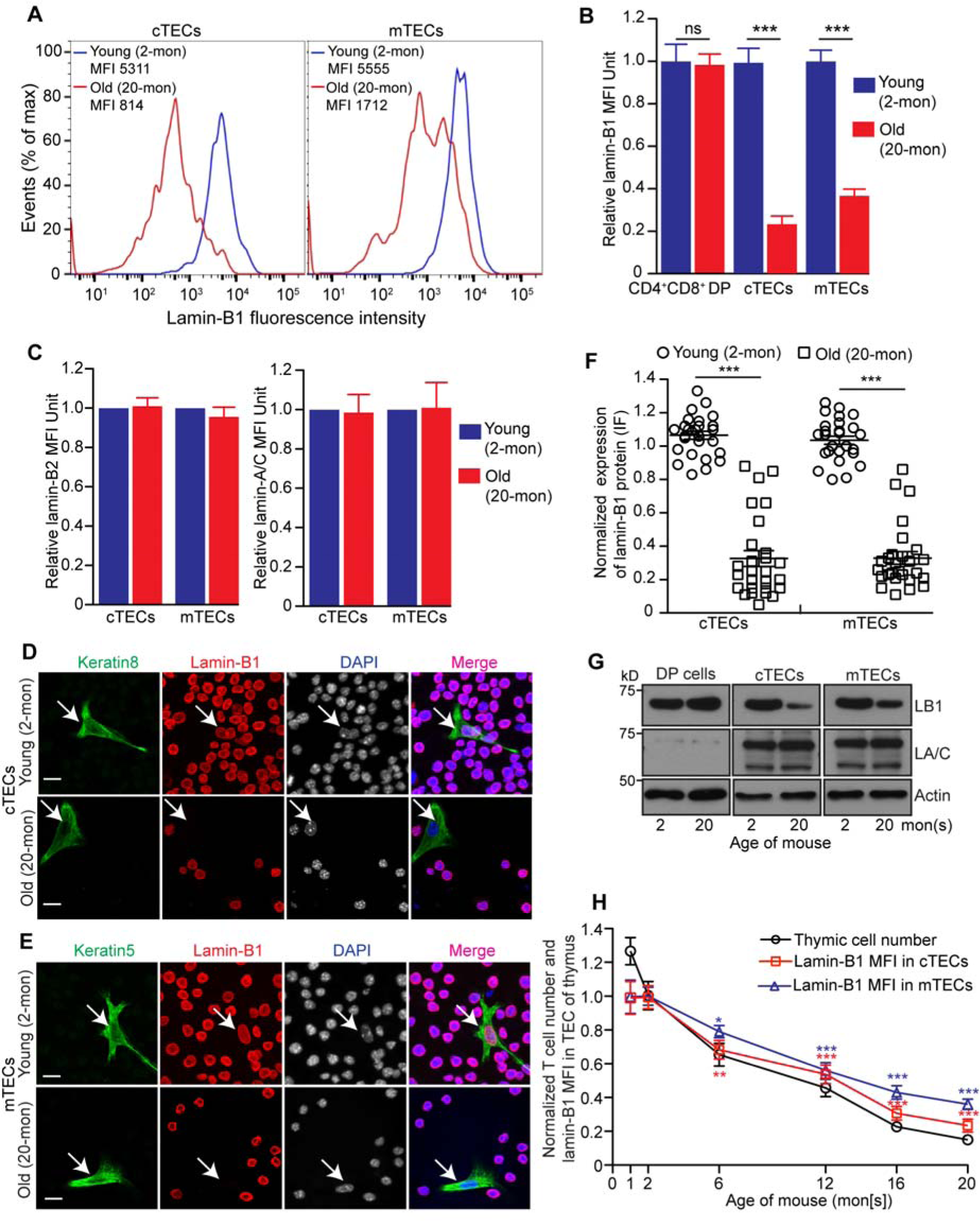
Effects of Aging on lamin-B1 protein levels in TECs. (A) Flow cytometry analyses of lamin-B1 levels in cTECs (left) and mTECs (right) from 2-(young, blue) or 20-mon-old (old, red) wild type mouse thymuses. Shown are representative histogram plots (out of >3 independent experiments) with the mean fluorescence intensities (MFI) of lamin-B1 indicated. (B) Quantification of lamin-B1 MFI in the CD4^+^CD8^+^ double-positive (DP) thymocytes, cTECs, and mTECs in the young and old WT mouse thymuses. (C) Quantification of lamin-B2 (left) and lamin-A/C (right) MFI in the cTECs and mTECs in the young and old WT mouse thymuses from two mice analyzed on different days. (D-E) Immunostaining of lamin-B1 (red) in cTECs (D) and mTECs (E) along with the spiked-in RAW264.7 cells (ATCC, #TIB-71) as an internal control for lamin-B1 staining intensity. Keratin 8 or 5 (green) labels cTECs (D) or mTECs (E) isolated from 2- or 20-mon WT mouse thymuses, respectively. (F) Quantification of lamin-B1 fluorescence intensity in cTECs or mTECs from (D) and (E). The average lamin-B1 fluorescence intensity in TECs and the surrounding spiked-in RAW264.7 cells were measured and the TEC lamin-B1 fluorescence intensities were plotted relative to the lamin-B1 intensity in spiked-in RAW264.7 cells which was set to 1. 25 cTECs or mTECs were measured in three independent experiments. (G) Western blotting analyses of lamin-B1 and lamin-A/C in CD4^+^CD8^+^ DP cells, cTECs, or mTECs in 2- or 20-mon thymuses. One 2-mon-old or five pooled 20-mon-old thymuses were used for each Western blotting analysis. β-actin is used for loading control. Shown are one representative Western blots of two independent experiments. (H) Quantification of the total thymic cell numbers (black) and lamin-B1 MFI in cTECs (red) or mTECs (blue) at the indicated ages. Total thymic cell number or MFI of lamin-B1 in cTECs or mTECs were plotted relative to those of 2-mon WT thymus, which was defined as 1 (≥three independent experiments). Error bars, SEM. Scale bars, 20 μm. Student’s t test: *P<0.05, **P< 0.01, ***P<0.001, ns: not significant.

We next analyzed how the kinetics of lamin-B1 reduction in TECs correlated with the two phases of thymic involution. We quantified the total thymic cell number as an indicator for thymic involution and used FACS to determine lamin-B1 level by MFI in all TECs. We found that mouse thymuses exhibited ~20-25% reduced of total cell numbers within the first two 2 months after birth, but lamin-B1 in TECs remained unchanged in this time window, whereas a gradual reduction of lamin-B1 in TECs occurred and it coincided with the second phase of gradual thymic aging as judged by total thymic cellularity (Figure 3H). Thus lamin-B1 reduction correlates with the age-associated thymic involution.

### Lamin-B1 decline in TECs accompanies age-associated thymic inflammation and can be triggered by proinflammatory cytokines

Age-associated increase in intrathymic lipotoxic danger signals have been shown to lead to caspase-1 activation via the Nlrp3 inflammasomes in thymic macrophages, which can in turn promote thymic inflammation and age-related thymic demise (Youm et al., 2012), but the cellular and molecular targets of this inflammation in thymus is unknown. We first examined whether the elevated inflammatory state could correlate with age-related thymic involution by applying FACS to isolate macrophages in thymuses dissected from 2, 6, 12, 16, and 20-mon-old mice (Figure 4A). Reverse transcriptase quantitative polymerase chain reaction (RT-qPCR) and Western blotting analyses were used to measure the key proinflammatory cytokines, TNF-α, IL-1β, IL-1α, and IL-6, that have been implicated in thymic degeneration (Billard et al., 2011). We found that the thymic macrophages exhibited a significantly increased expression of proinflammatory cytokines by 6 months of age and the levels continued to increase with aging (Figure 4B and C). Similar to the thymic macrophages, we found that the thymic Sirpα^+^ dendritic cells (DCs) (Ki et al., 2014) also exhibited an increased expression of these proinflammatory cytokines (Figure S3).

**Figure 4.**
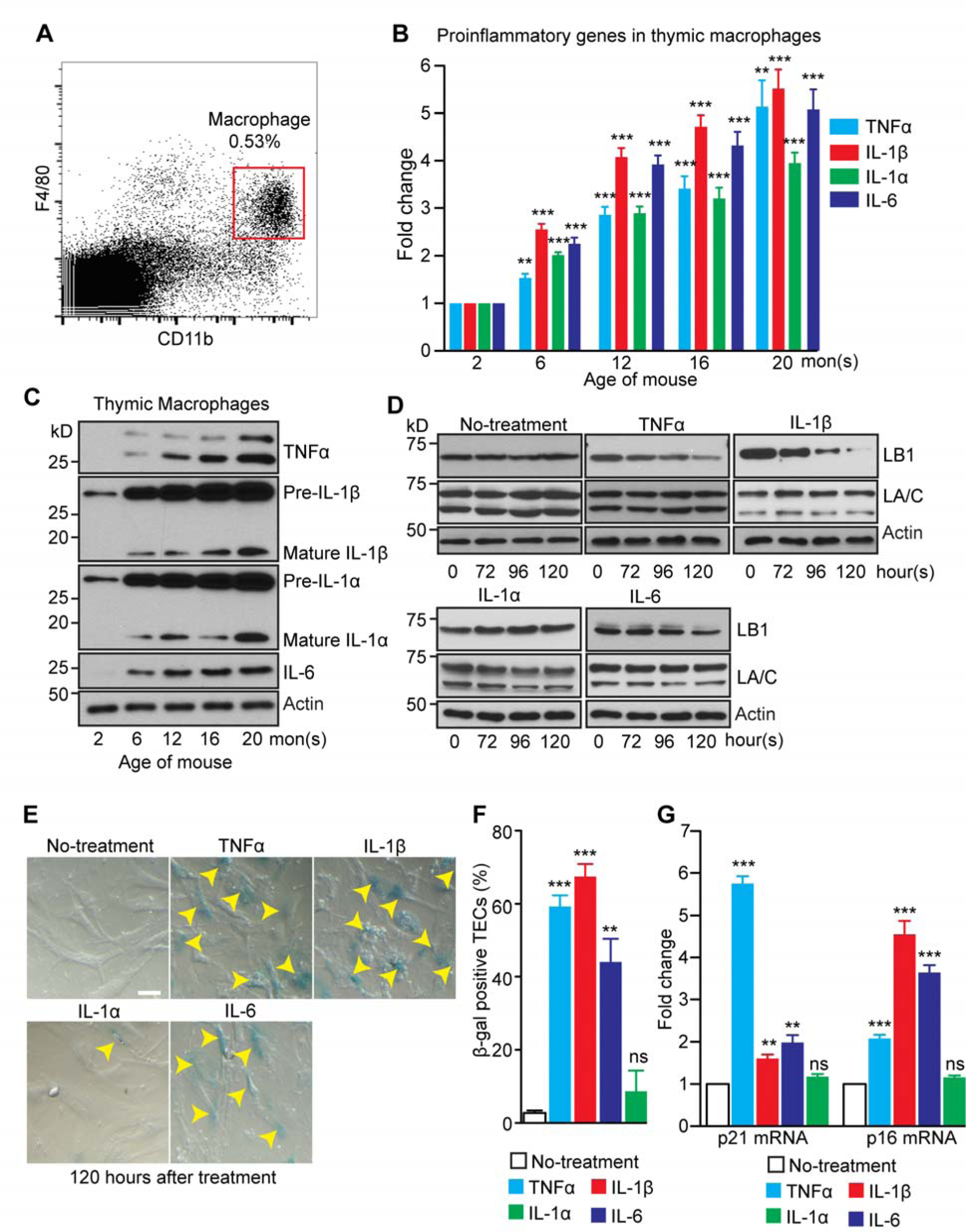
Effects of inflammatory cytokines on the TEC lamin-B1 level. (A) The flow cytometry gating strategy for sorting macrophages from a 2-mon-old WT thymus. Macrophages were identified based on the expression of F4/80 and CD11b. (B) RT-qPCR analyses of TNF-α, IL-1β, IL-1α, and IL-6 in macrophages isolated from WT thymuses at the indicated ages. The increased expression was plotted relative to the 2-mon-old thymus, which was set to 1. (C) A representative Western blotting analysis of TNF-α, IL-1β, IL-1α, and IL-6 in macrophages isolated from WT thymuses at the indicated ages. β-actin, loading control. (D) Western blotting analyses of lamin-B1 and lamin-A/C in cultured primary TECs treated with TNF-α (10 ng/ml), IL-1β (20 ng/ml), IL-1α (20 ng/ml), and IL-6 (20 ng/ml) for the indicated hours. (E) Increased TEC senescence as judged by /-galactosidase staining (yellow arrowheads) upon treatment by TNF-α, IL-1β, and IL-6, but not by IL-1L. (F) Quantification of the percentage of β-galactosidase positive primary TECs. Total 150 cells from 3 biological replicates were counted for each experiment. (G) RT-qPCR analyses showing the up regulation of p21 and p16 in the primary TECs upon treatment with the indicated proinflammatory cytokines. Scale bar, 20 ym. Error bars, SEM from three biological repeats. Student’s t test: *P<0.05, **P<0.01, ***P<0.001, ns: not significant. See also Figure S3 and S4.

Since by 6 months of age, the lamin-B1 level in TECs was reduced by ~30% (see Figure 3H), we reasoned that the increased proinflammatory cytokines detected in thymus at this age could contribute to lamin-B1 reduction in TECs. To test this, we isolated and cultured primary TECs from 2-mon-old mouse thymuses and treated these cells with TNF-α, IL-1β, IL-1α, and IL-6. Whereas TNF-α, IL-1β, and IL-6 induced a gradual reduction of lamin-B1 without affecting lamin-A/C, IL-1α had no effect on either lamins (Figure 4D). This shows that TEC lamin-B1 levels are sensitive to certain proinflammatory cytokines. Since proinflammatory cytokines are known to trigger various signaling pathways to initiate and maintain cellular senescence (Acosta et al., 2008; Kuilman et al., 2008) and since the senescence of human and mouse fibroblasts is accompanied by lamin-B1 reduction (Coppe et al., 2008; Dreesen, Chojnowski, et al., 2013), we examined whether TNF-α, IL-1β, and IL-6 triggered TEC senescence. By quantifying cells positive for, β-galactosidase, a senescence marker, we found that >40% of TECs underwent senescence after treatment by TNF-α, IL-1β and IL-6, but not by IL-1α (Figure 4E and F). Consistently, RT-qPCR analyses of p21 and p16, two cell cycle inhibitors involved in p53- and retinoblastoma protein (Rb)-mediated senescence, respectively (Rodier & Campisi, 2011), showed that TNF-α induced a strong up-regulation of p21 expression while IL-1β and IL-6 mainly engaged in inducing p16 (Figure 4G) These findings suggest that the age-associated increase of thymic TNF-α, IL-1β, and IL-6 can trigger lamin-B1 reduction in TECs at least in part by inducing cellular senescence.

Thymic involution also occurs under pathophysiological conditions, including chronic and acute infection (Chinn et al., 2012). We tested whether lamin-B1 reduction in TECs could be stimulated by cytokines via injecting lipopolysaccharide (LPS), which is known to cause endotoxemia and acute thymic atrophy (Billard et al., 2011). Consistent with previous report, intraperitoneal injection (IP) of one-dose of LPS (100 μg) (Figure S4A) induced acute thymic involution within 48 hours as measured by total thymic cell number counts (Figure S4B). By 48 hours post LPS injection, we found a significant reduction of lamin-B1 levels in FACS-sorted mTECs by Western blotting analyses, while lamin-A/C did not exhibit obvious changes (Figure S4C). Due to a very small number of cTECs present in the young thymuses, we were unable to assess lamin protein levels in these cells. Taken together these analyses suggest that age-associated increase in the proinflammatory cytokines TNF-α, IL-1β, and IL-6 can induce TEC senescence and lamin-B1 reduction.

### Lamin-B1 is required in TECs to support adult thymus organization in part by maintaining proper gene expression

The role of TEC lamin-B1 in thymus development prompted us to ask whether during adulthood lamin-B1 is required for maintaining thymus structure and function, and if the age-associated lamin-B1 reduction in TECs contributes to thymic involution. We employed *K8* or *K5CreER^T2^* to delete *Lmnb1* in either cTECs or mTECs, respectively, in the adult thymus by injecting tamoxifen (TAM) starting at 2 months of age (Figure 5A) (Cheng et al., 2010). It is important to note that cTEC- and mTEC-restrict lineages are initially derived from biopotent TEC progenitors that may express both K5 and K8 (Klug et al., 1998). Thus, *K5CreER*^*T2*^ may delete *Lmnb1* alleles in some cTEC sub-lineages, whereas *K8CreER*^*T2*^ may delete *Lmnb1* alleles in some mTEC sub-lineages. Since TAM-induced genomic excision have variable efficiency depending on the TAM dosages, *CreER*^*T2*^ lines, and specific tissues, we tested different TAM doses. We found that 1mg TAM/10g body weight/day for five consecutive days followed by feeding the mice drinking water containing TAM (25µg/ml) resulted in an obvious reduction in thymic sizes in the *Lmnb1^f/f^;K8CreERT2* or *Lmnb1*^f/f^*;K5CreER*^*T2*^ mice compared to the littermate controls and this procedure did not cause pronounced toxicity in all the injected mice. qRT-PCR revealed that this TAM regimen resulted in ~52% or 41% reduction of *Lmnb1* mRNA in the FACS-sorted cTECs or mTECs, respectively, at one month after the last TAM injection (Figure 5B). The size and total cell number of thymuses in *Lmnb1^f/f^;K8CreERT2* or *Lmnb1*^f/f^*;K5CreER*^*T2*^ mice were also reduced to 50-60% of the littermate controls (Figure 5C-E). These findings demonstrate that lamin-B1 reduction in either cTECs or mTECs in the adult thymus can cause thymic involution.

**Figure 5.**
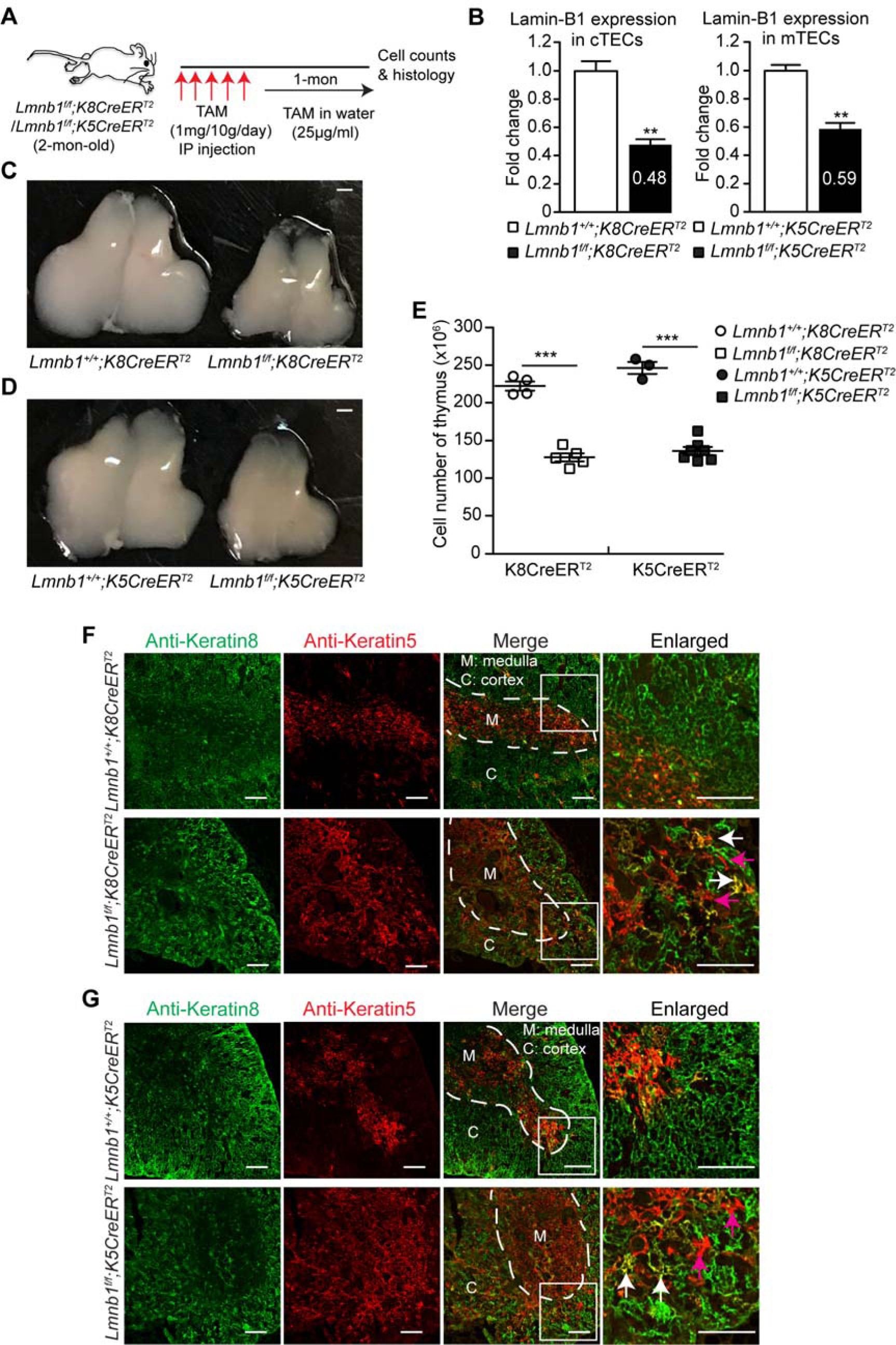
Effects of postnatal lamin-B1 deficiency in TECs on thymus size and organization. (A) The tamoxifen (TAM)-mediated *Lmnb1* deletion scheme. Keratin 8 (*K8*)- or keratin 5 (*K5*)-driven *CreER*^*T2*^ was used to delete *Lmnb1* in cTECs or mTECs, respectively, by injecting TAM followed by feeding in TAM-containing drinking water before analyses. (B) RT-qPCR analyses of lamin-B1 mRNA levels revealing the *Lmnb1* deletion efficiency by *K8CreER*^*T2*^ in cTECs (left) or by *K5CreER*^*T2*^ in mTECs (right). The average mRNA levels of lamin-B1 were calculated from 3 independent experiments with the control set to 1. (C-D) Representative images of thymuses from mice with indicated genotypes after TAM treatment as shown in (A). Scale bars, 1 mm. (E) The total thymic cells were counted and plotted for indicated genotypes as shown. (F-G) Keratin staining of thymuses from mice with indicated genotypes treated by TAM as shown in (A). White dash lines demarcate the cortical and medulla junction regions (CMJ) in the thymus. A section (white squares) of the thymus in each genotype was enlarged to show the well-defined (in the controls) or disorganized (in the mutants) CMJ. Arrows mark the spreading of K5^+^K8^+^ TECs (white) and K5^+^ TECs (purple) into the cortical regions in the in *Lmn1^f/f^;K8CreERT2* (F) or *Lmn1^f/f^;K5CreERT2* (G) thymuses. Scale bars, 100 m. Error bars, SEM from ≥3 independent experiments. Student’s t test: *P<0.05, **P< 0.01, ***P<0.001, ns: not significant. See also Figure S5.

We next performed K5 and K8 immunostaining to determine if lamin-B1 reduction affected the structural organization of the thymus. Compared to the littermate controls, the cortical and medullary compartments were disrupted with the spreading of K5^+^ mTECs into the cortical regions in the thymuses of TAM-treated *Lmnb1^f/f^;K8CreERT2* (Figure 5F) and *Lmnb1^f/f^;K5CreERT2* mice (Figure 5G). Our data demonstrate that lamin-B1 is required in the adult cTECs and mTECs to maintain the proper segregation of the cortical and medullary thymus architecture. Thus, the age-associated lamin-B1 reduction in TECs can contribute to the age-associated phase of thymic involution.

To explore further how lamin-B1 reduction contributes to thymic involution, we performed RNA-seq of the control and lamin-B1-depleted TECs. Since the deletion efficiency of *Lmnb1* in mTECs by *K5CreER*^*T2*^ is low and since the total mTEC number is also low in the TAM treated *Lmnb1^f/f^;K5CreERT2* thymus (Figure 5B), a high percentage of our isolated mTECs were WT for *Lmnb1* which makes it difficult to interpret the RNA-seq results. Therefore, we focused our analyses on the effect of lamin-B1 deletion in cTECs. We considered the differentially expression of genes as those with fold changes ≥ 2 than the control cTECs with the adjusted false discovery rate (FDR) <0.05 and P values <0.05 (see the method section). We found lamin-B1 reduction in cTECs resulted in the transcriptional up- and down-regulation of 533 and 778 genes, respectively (Figure S5A; supplement table 1 for the full list). DAVID Gene Ontology (GO) analysis revealed that lamin-B1 deletion resulted in a dis-regulation of genes implicated in cell adhesion, immune system process and development, T cell differentiation, and cytokine production that are relevant to thymus function (Figure S5B-D; supplement table 2 for the full list). Thus, lamin-B1 plays an important role in the expression of genes required for maintaining TEC structure and niche, which would in turn support T cell development.

### Lamin-B1 reduction in the adult TECs decreases thymopoiesis and accelerates age-related loss of naïve T cells

Since the intact TEC compartments play essential roles in thymopoiesis, we next investigated the impact of lamin-B1 reduction in the adult TECs on T-cell development. We analyzed the CD4^+^ SP and CD8^+^ SP thymocytes and found a ~40-45% reduction of these SP thymocytes in the *Lmnb1^f/f^;K8CreERT2* mice compared to littermate controls one month after the last TAM injection (Figure S6A-C), indicating that lamin-B1 plays an important role in cTECs to support efficient thymopoiesis in the adult mouse thymus. We also observed a similar but milder effect upon lamin-B1 reduction in the adult mTECs in the *Lmnb1^f/f^;K5CreERT2* mice (Data not shown), which is consistent with the less efficient *Lmnb1* deletion in these mice than that of the *Lmnb1^f/f^;K8CreERT2* mice. Thus lamin-B1 reduction upon aging can contribute to the gradual decline of the production of naïve T-cells in the thymus.

Since the decline of naïve T cells in the peripheral immune system upon thymus involution contributes to the aging of the immune system (often referred to as immunosenescence) and reduced immune surveillance in elderly populations (Youm et al., 2012), we next assessed whether TAM-induced lamin-B1 deletion in the adult cTECs in thymus could affect the CD4^+^ and CD8^+^ T-cell compartments in the peripheral immune organs such as the spleen and the mesenteric lymph node (mLN). Although no changes occurred in the frequency and the number of CD4^+^ and CD8^+^ naïve T cells (CD62L^+^CD44^-^) in these two immune organs by one month after the last TAM injection, by 6 months, we found a pronounced reduction of the naïve T cells in the *Lmnb1^f/f^;K8CreERT2* mice compared to their littermate controls (Figure S6DI). This reduction was accompanied by a marked increase of CD4^+^ and CD8^+^ effector memory T cells (EM, CD62L^-^CD44^+^) and CD8^+^ central memory T cells (CM, CD62L^+^CD44^+^) in these organs upon lamin-B1 reduction (Figure S6D and G). Together, these findings demonstrate that lamin-B1 reduction in adult TECs promotes thymic involution and causes changes in the peripheral T cell composition that are reminiscent of the age-associated alterations in the peripheral immune system.

### Identification of new adult TEC subsets and the role of lamin-B1 in maintaining these subsets in the thymus

To understand how natural aging and postnatal lamin-B1 reduction contribute to TEC maintenance, we performed droplet-based scRNA-seq on isolated total TECs (DAPI^-^CD45^-^ EpCAM^+^) from the WT young (2-mon), WT old (20-mon), and TAM-treated 3 mon-old *Lmnb1^f/f^;K8CreERT2* mouse thymuses (see Figure 5A for the TAM scheme). For the reasons mentioned above in transcriptome analyses of population TECs, we focused on using *K8CreER*^*T2*^ for postnatal lamin-B1 deletion in scRNA-seq. Due to the expression of both K5 and K8 in bipotent TEC progenitors, *K8CreER*^*T2*^ should delete *Lmnb1* in all cTECs and a subset of mTECs.

After quality control, we retained 5,872 cells with 2,149 median genes per cell from the young, 6,973 cells with 1,948 median genes per cell from the old, and 7,314 cells with 2,416 median genes per cell from the TAM-treated *Lmnb1^f/f^;K8CreERT2* thymus. We combined all three samples to assess TEC heterogeneity. This also allowed us to assess how aging and postnatal lamin-B1-depletion affect TEC composition. Unsupervised clustering analysis by t-stochastic neighborhood embedding (tSNE) identified 17 major clusters (C1-17) of TECs in our dataset (Figure 6A and B; supplement table 3 for the full list). We found that the 17 clusters are present in all three thymus samples (Figure 6C-E), suggesting that our profiling identified the major TEC subsets in adult mouse thymus.

**Figure 6.**
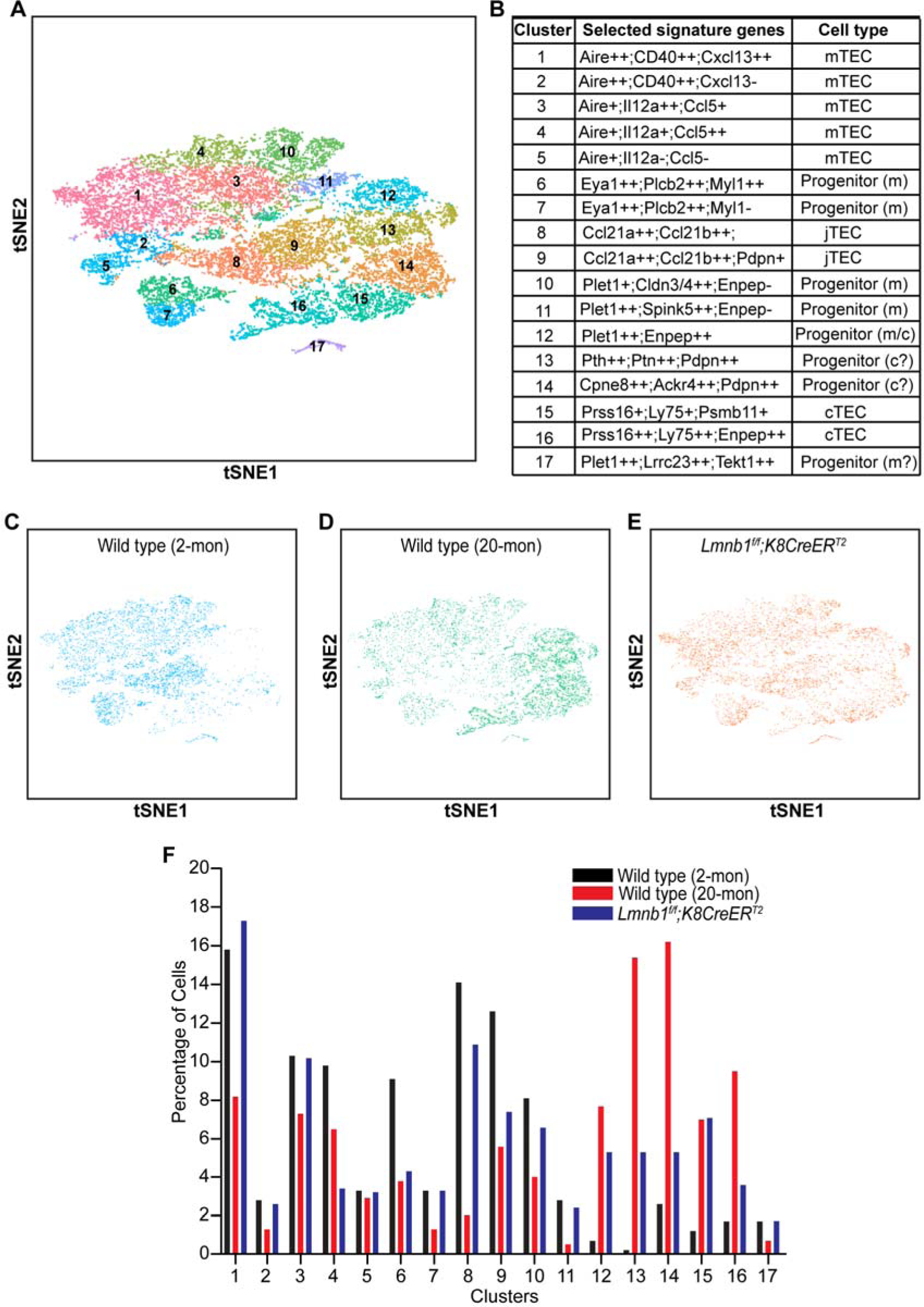
Single-cell RNA-seq of TECs in young, old, and postnatal lamin-B1 depleted thymus. (A) t-distributed Stochastic Neighbor Embedding (tSNE) plot of combined TECs from WT young (2-mon; 5872 cells), WT old (20-mon; 6973 cells), and TAM-treated *Lmnb1^f/f^;K8CreERT2* (7314 cells) mouse thymuses. Cell clusters are assigned to specific subgroups (1-17) based on differentially expressed marker genes. Each point is a single cell colored by cluster assignment. (B) List of selected marker genes and predicted subpopulations of individual TEC clusters. +: 2≤ fold change <3; ++: fold change ≥3; -: not enriched. (C-E) tSNE plots of TECs derived from WT young (C), WT old (D), and TAM-treated *Lmnb1^f/f^;K8CreERT2* (E) mouse thymuses. Each point is a single cell colored by cluster assignment. (F) Percentage of the 17 TEC clusters shown in B in WT young, WT old, and TAM-treated *Lmnb1^f/f^;K8CreERT2* mice See also Table S3.

Based on the enriched mTEC signature genes such as Aire (‘Aire regulates negative selection of organ-specific T cells,’; M. S. Anderson et al., 2002), the cell clusters C1-5 represent five differentiated mTEC subsets (Figure 6A-B). The cell clusters C6 and C7 do not appear to express the well-characterized mTEC signature genes. However, when analyzing the available transcriptome profiles of the mouse neonatal cTECs and mTECs (St-Pierre et al., 2013), we found that C6 and C7 express high levels of genes, such as Plcb2 and Myl1, that are specifically expressed in mTECs but not in cTECs in the neonatal mouse thymus. More importantly, these two clusters are enriched in expression of Eya1, a gene specifically expressed in a subset of mTEC progenitors identified by a recent scRNA-seq of embryonic and neonatal thymus (Kernfeld et al., 2018). Thus, C6 and C7 may represent two previously unappreciated mTEC progenitor subsets in adult mouse thymus (Figure 6B).

We also identified several cell clusters, C8-C11, that express different genes previously reported to be expressed in mTEC progenitors. Ccl21a and Ccl21b have been reported as markers for junctional TEC precursor (jTEC) that could give rise to fully differentiated mTECs (Miragaia et al., 2018; St-Pierre et al., 2013). We found the C8 and C9 clusters express these two jTEC markers, but they can be further separated from one another based on differential expression of other genes such as Pdpn (Figure 6B), suggesting that the previously identified jTECs can be divided into two subsets. The C10 and C11 clusters represent two additional subsets of mTEC-restricted progenitors because they express TEC progenitor specific marker Plet1 but lack the expression of Enpep, which encodes the antigen epitope for Ly51 (Figure 6B), one of the markers for bipotent TEC progenitor (Depreter et al., 2008; Ulyanchenko et al., 2016). By contrast, the cluster C12 expresses both Plet1 and Enpep and it corresponds to the newly identified Plet1^+^Ly51^+^ bipotent TEC progenitor that can effectively generate both cTECs and mTECs in the adult mouse thymus (Ulyanchenko et al., 2016). Finally, a minor cluster C17 may belong to the mTEC-restricted progenitors because these cells express Plet1 and are enriched for genes specifically expressed in mTECs, such as Lrrc23 and Tekt1 (St-Pierre et al., 2013).

The scRNA-seq also revealed two different adult cTEC subsets as cluster C15 and C16 because they share genes previous identified for cTECs and they can be separated by some previously unappreciated differentially expressed genes (Figure 6B). Interestingly, we identified two clusters C13 and C14 that do not express previously defined mature cTEC signature genes, such as Prss16 and Ly75. However, their expression of specific genes enriched in cTECs but not in mTECs in neonatal thymus, including Pth, Ptn, Cpne8 and Ackr4 (St-Pierre et al., 2013), suggests that they represent two new cTEC-restricted short-term progenitors. Together, our scRNA-seq profiling has revealed new TEC progenitors and mature mTEC and cTEC subsets in adult thymus.

The identification of different TECs provides us the opportunity to assess how natural aging affects TEC composition changes and whether lamin-B1 reduction in adulthood contributes to ageing-associated TEC composition changes. We first examined the frequency of individual TEC subsets in WT young and old mouse thymuses. Consistent with previous phenotypic analyses showing an overall reduction of mTECs upon aging (Chinn et al., 2012), we found that the percentages of most mTEC subsets (C1-4) were significantly reduced upon aging (Figure 6F). Importantly, we identified a unique group of mTECs belonging to C5 that remained unchanged upon aging. By contrast, the two mature cTEC subsets (C15 and C16) exhibited a strong increase in percentage upon aging, which is consistent with previous finding that total cTEC numbers increase with increased age (Chinn et al., 2012).

Of the progenitor cells identified, all the mTEC progenitor subsets (C6-11) and the minor mTEC progenitor C17 exhibited a clear reduction upon aging (Figure 6F). Unexpectedly, we found that the bipotent TEC progenitor cluster C12 is present at a much higher frequency in the aged thymus than that of the young (Figure 6F). This suggests that the bipotent TECs can provide the source of thymic regeneration in aging mice.

When analyzing the impact of forced postnatal lamin-B1 depletion by TAM treatment of *Lmnb1^f/f^;K8CreERT2* mice, we found an increase in all cTEC progenitor and mature cTEC subsets similar to those observed upon natural thymic aging (Fig. 6D-F). Similar to the WT old thymuses, postnatal lamin-B1 depletion in TAM-treated *Lmnb1^f/f^;K8CreERT2* thymus resulted in a reduction of C4 mTEC subset (Figure 6F). We also noticed a mild reduction of mTEC-restricted TEC progenitors (C6, C8-10) in TAM-treated *Lmnb1^f/f^;K8CreERT2* thymus (Figure 6F). These indicate that *K8CreER*^*T2*^ can delete *Lmnb1* alleles in some mTECs due to K8 expression in certain progenitor cells. Additionally, both the naturally aged WT thymus and the postnatal lamin-B1 depleted thymus exhibited a similar increase in the bipotent TEC progenitor subset C12 (Fig. 6B and F). Taken together sc-RNA-seq analyses demonstrate that postnatal lamin-B1 reduction contributes to the age-associated changes of TEC composition, thereby leading to thymic involution and disfunction.

## Discussion

Since organ building and maintenance require structural proteins that support cell and tissue shapes, it is natural to assume that aging-associated deterioration of these proteins would contribute to organ degeneration. The complex cellular composition in each organ, the difference in organ function, and the varying rates of organ aging have, however, made it challenging to identify structural proteins that have cell type specific functions in both building and maintaining organs. It is also difficult to define the causes and consequences of structural protein disfunction in the same cell type later in life that contributes to organ degeneration. Our studies of the thymus demonstrate that among the three lamins found in mammals, lamin-B1 is specifically required in TECs for thymus development and maintenance. Our findings further suggest that the age-associated increase in intra-thymic inflammation triggers senescence-associated lamin-B1 reduction in TECs, which promotes the gradual thymic degeneration and dysfunction.

Cellular senescence has been implicated in promoting organ aging (Campisi, 2013). However, senescence triggers multitudes of cellular changes, which make it difficult to tease apart the major and minor impact of these changes on organ aging. Additionally, although extensive studies of tissue culture cells have shown that many perturbations, including oxidative stress, DNA damage, oncogene transformation, and inflammation, can induce senescence *in vitro* (Ren et al., 2009), which cell type in a given organ is most sensitive to these senescence triggers and whether a specific senescence change of a given cell type makes a key contribution to organ aging remains unknown. By using mouse thymus known to experience age-associated activation of inflammasomes (Youm et al., 2012), our studies provide a connection between gradual increase of thymic inflammation to TEC senescence and lamin-B1 reduction.

We show that both thymic macrophages and Sirpα^+^ DCs exhibit age-associated increase in IL1-α, IL-1β, IL-6, and TNF-α, Among these proinflammatory cytokines, we found that IL-1β, IL-6, and TNF-α induced senescence and lamin-B1 reduction in TECs. In addition, we observed that TEC lamin-B1 was reduced upon acute induction of thymic involution by inflammatory challenges. These findings suggest that both chronic and acute thymic inflammation targets TECs by inducing their senescence and lamin-B1 reduction. Since lamin-B1 reduction in TECs and the elevation of IL1-α IL-1β, IL-6, and TNF-α in the thymic myeloid cells occur 2 months after birth, these thymic changes do not contribute to the first developmental phase of thymic involution but are correlated with the age-associated thymic degeneration.

Previous studies have shown that *Foxn1*, whose expression is essential for TEC proliferation and differentiation, undergoes an age-associated decline in TECs. Since Foxn1 deficiency in mouse TECs triggers premature thymic atrophy, the postnatal down-regulation of Foxn1 has been generally accepted as a major mediator of age-associated thymic involution (L. Chen et al., 2009; Cheng et al., 2010). More recently, however, it is recognized that mouse thymus displays a developmental and age-associated phases of involution (Aw & Palmer, 2012; Shanley et al., 2009). The developmental involution occurs ~6 weeks after birth when thymus has produced a large number of naïve T-cell required for establishing the peripheral immune system. Therefore, this initial thymic reduction could help to divert resources for other bodily needs without compromising the immune functions. On the other hand, the age-associated involution is more gradual, and as thymic atrophy progresses, the immune system is negatively affected by the significant reduction of naïve T cell production. The differences in the two phases of thymic involution lead to the idea that they may be controlled by different mechanisms (Shanley et al., 2009). Since the accumulation of Foxn1 negative (Foxn1^-^) TECs reaches a plateau in ~50% of the total TECs within 10 weeks after birth (O’Neill et al., 2016; Rode et al., 2015), Foxn1 reduction soon after birth appears to be responsible for the developmental thymic reduction. Consistent with this, overexpression of *Foxn1* in TECs can delay, but not prevent the age-dependent involution (Zook et al., 2011). Therefore, it is critical to identify additional TEC intrinsic factors that contribute to age-dependent phase of involution. Our studies suggest that intra-thymic inflammation can lead to TEC senescence and lamin-B1 reduction. Lamin-B1 reduction in turn contributes to the age-dependent thymic involution.

Using germline deletion of *Lmnb1* or *FN1Cre*-mediated *Lmnb1* deletion in TECs during embryogenesis, we show that lamin-B1 is required for the proper building of the cortical and medulla TEC compartments. This is consistent with a known function of B-type lamins in tissue building. By carefully characterizing the roles of lamins in the thymus, we reveal a specific requirement of lamin-B1 in TECs but not in the T-cell lineage. Indeed, we found that lamin-B1 helps to balance proper TEC differentiation from a common bipotent TEC progenitor and mediates terminal differentiation of cTECs as *Lmnb1*-deficient thymus exhibits skewing towards the immature MHCII^low^cTEC dominant compartment.

To test if lamin-B1 reduction after birth could contribute to age-associated thymic involution, we induced *Lmnb1* deletion in TECs starting from 2 months after birth and found it advanced thymic degeneration and dysfunction as judged by an increased disorganization of the TEC compartments, decreased naïve T cell production and lymphopenia.

By applying sc-RNA-seq to TECs isolated from young WT, old WT, and postnatal lamin-B1 depleted thymuses, we identified 17 subsets of TEC progenitors and differentiated mTECs and cTECs in adult thymus. The signature genes we identified in different TEC subsets can be used as biomarkers for isolation and functional studies of specific TECs both *in vitro* and *in vivo*. The newly defined adult TEC subsets in this study have allowed us to explore in depth about how aging and postnatal lamin-B1 depletion in TECs affect TEC composition changes. Importantly, we found an overall similar change of composition in TEC subsets in TAM-treated *Lmnb1^f/f^;K8CreERT2* and WT aged mouse thymuses compared to that of the young. Specifically, we uncovered one mTEC progenitor subset that do not appear to change and two bipotent TEC progenitor subsets that increase upon aging or lamin-B1 depletion. Together, these findings strongly suggest that lamin-B1 reduction in TECs upon aging plays a key role in triggering the age-associated alteration of TEC composition, thus contributing to thymic involution. The newly defined adult TEC subsets should also pave the way for further dissecting the functional contributions of individual TEC subsets to thymic aging and regeneration.

Additionally, we show that postnatal lamin-B1 reduction contributes to significant transcriptome changes enriched for genes that function in cell proliferation, differentiation, cell-cell adhesion, and cytokine production. These gene expression changes help to explain why lamin-B1 reduction upon aging or upon Cre-mediated gene deletion in TECs causes thymic tissue disorganization, reduced naïve T cell production, and lymphopenia.

Lamins are known to influence the expression of genes found both in the lamina-associated chromatin domains (LADs) and non-LADs. Our recent studies of lamin null mouse embryonic stem cells (mESCs) have shown that lamins regulate the 3D chromatin organization by ensuring the proper condensation and position of LADs at the nuclear periphery (X. Zheng et al., 2018). The decondensation and detachment of LADs upon lamin loss can lead to altered gene expression by bringing the active chromatin domains into the close vicinity of inactive ones. Since lamin-B reduction in *Drosophila* fat body cells and lamin-B1 reduction in TECs both cause expression changes of genes in LADs and outside of LADs, it would be interesting to determine if the gene expression changes in these cells are facilitated by 3D genome organization changes upon lamin-B depletion.

## Experimental Procedures

### Mice

All mouse experiments were approved by the Institutional Animal Care and Use committee of Carnegie Institution for Science. *Lmnb1*^*f/f*^ allele was derived from *Lmnb1*^tm1a(EUCOMM)Wtsi^ by the EUCOMM project. The conditional *Lmnb2* allele, *Lmnb2*^*f/f*^, was derived from *Lmnb2*^tm1a(KOMP)Wtsi^ (KOMP project) by breeding with ACTB-FLPe mice to remove neomycin cassette flanked by Frt sites. *Lmna*^f/f^ and *Lmnb1*^-/-^ alleles were generated in house by Dr. Youngjo Kim and were reported previously (Y. Kim & Zheng, 2013). All mice were further backcrossed to the C57BL/6 background for at least 6 generations. *LckCre*, *Foxn1Cre*, *K5CreET*^*T2*^, *K8CreER*^*T2*^ and *OT-II* mice were purchased from the Jackson Laboratory. *K5CreET*^*T2*^ and *K8CreER*^*T2*^ mice were further backcrossed to the C57BL/6 background for at least 2 generations. 20-mon-old female C57BL/6 mice were purchased from National Institution on Aging. Other mice in cohort experiments at the indicated ages were raised by the mouse facility of Carnegie Institution for Science. All animals were housed under a 12-/12-hour light dark cycle and fed *ad libitum*.

### Primary TEC isolation and culture

TECs were prepared according to a published protocol with modifications (Jain & Gray, 2014). In brief, thymic lobes were cut into 2 mm pieces and then treated for 15 min at 37°C with an enzymatic mixture containing 0.25 U Liberase TM (Roche) and 0.1% DNase I (Sigma-Aldrich) in RPMI 1640. The supernatant was collected, and the digestion was repeated twice. Cells were filtered through 100 μm cell strainer and spun at 1200 rpm for 5 min. The CD45^+^ cells were depleted by CD45 microbeads (Miltenyi Biotec 130-052-301) and the enriched CD45^-^ cells were then stained with DAPI, CD45, EpCAM, Ly51, I-A/I-E (Biolegend) and UEA-1 (Vector laboratories) for 20 min at 4°C followed by sorting with FACSAria^TM^ III (BD Bioscience). The TEC subpopulations were identified as, mTEC: CD45^-^EpCAM^+^UEA-1^+^Ly51-MHCII^+^; cTEC: CD45^-^EpCAM^+^UEA-1^-^Ly51^+^MHCII^+^. FACS sorted TECs were further subjected to either downstream analyses or culture. For primary TEC culture, ~50,000 TECs were seeded in 48-well culture plates and cultured for up to 7 days in Dulbecco’s modified Eagle’s medium nutrient F12 (Invitrogen) supplemented with 3 μg/ml insulin (Sigma-Aldrich), 20 ng/ml epidermal growth factor (PeproTech), 100 unit/ml penicillin-streptomycin, and 10% fetal bovine serum. Cultures were maintained at 37°C and 5% CO2, and the medium was changed every two days. For pro-inflammatory cytokine treatment, TECs were cultured with above-mentioned medium supplemented with different concentrations of cytokines (10 ng/ml TNF-α, PeproTech; 20 ng/ml IL-1β, Biolegend; 20 ng/ml IL-6, PeproTech; 20 ng/ml IL-1α, Biolegend). TECs were collected at indicated time points for RNA or protein extraction.

### Flow cytometry analysis of nuclear lamins

Single-cell suspension of freshly dissected thymus was prepared as described for TEC isolation without depletion of CD45^+^ cells. Cells were first treated with TruStain fcX kit (Biolegend) to block CD16/32 and Zombie UV™ fixable viability kit (Biolegend) to exclude dead cells before cell surface marker staining. Surface marker staining was used to distinguish different cell types within the thymus: CD4/CD8 (T cell subsets); mTEC: CD45^-^EpCAM^+^UEA-1^+^Ly51^-^; cTEC: CD45^-^EpCAM^+^UEA-1^-^Ly51^+^. After surface marker labeling, staining of lamins follows the detailed protocol described in the True-Nuclear Transcription Factor kit (Biolegend). Alexa Fluro 647 antibody labeling kit (Life technology) was used to prepare fluorochrome-conjugated lamin antibody. In brief, 50 μg rabbit anti-lamin-B1 (Abcam), mouse anti-lamin-B2 (Invitrogen, E3), mouse anti-lamin-A/C (Active Motif) and control rabbit or mouse IgG (Santa Cruz) was labeled following the manufacturer’s protocol. Fluorochrome-conjugated antibodies were reconstituted at 0.5 mg/mL in PBS and 0.1 μg of each antibody was used for staining 10^6^ cells in 100 μl volume. All samples were stained for 20 min at 4°C if not specifically indicated and then were immediately processed for FACS analyses by FACSAriaTM III (BD Bioscience). Data was analyzed with FlowJo software (Tree Star).

### Flow cytometry analysis and sorting

Single-cell suspension of the thymus was prepared by passing minced tissue through a 40 μm cell strainer. Cells were treated with TruStain fcX kit (Biolegend) to block CD16/32 before staining. All surface markers were stained for 20 min at 4°C, if not specifically indicated, and then were immediately processed for FACS analyses or sorting by FACSAriaTM III (BD Bioscience). The following FACS monoclonal antibodies were used for experiments: CD45, EpCAM, Ly51, I-A/I-E (MHCII), CD4, CD8, TCRβ, TCRγδ, CD69, TCRβV5.1, CD11b, F4/80, CD11c, CD172 (Sirpα) (all from Biolegend) and UEA-1 (Vector laboratory). Macrophage and DC subsets were directly sorted into Trizol (Life technology) for RNA or 4X Laemmli sample buffer (Bio-rad) for protein extraction. Data was analyzed with FlowJo software (Tree Star).

### Immunofluorescence staining

Freshly isolated thymuses were embedded in OCT (Leica microsystems) and frozen immediately in −80°C. For keratin staining, 10 μm sections were fixed in a 1:1 mixture of acetone and methanol at −20°C for 10 min and then washed with PBS for 3 times. After blocking, samples were stained with primary antibodies containing rat-anti-keratin 8 (Troma-1, DSHB, 1:20) and rabbit-anti-keratin 5 (Biolegend, 1:300) at room temperature (RT) for 1 hour, followed by Alexa488 goat anti-rat and Alexa568 goat anti-rabbit (Invitrogen) secondary antibodies. Samples were then mounted with ProLong^®^ Gold Antifade (Fisher Scientific) and allowed to dry overnight at RT before imaging. For lamin-B1 immunostaining (IF) assay, cTECs and mTECs were collected by FACS sorting and then attached to glass slides with spike-in RAW264.7 cells by cytospin (Shandon Cytospin 4) spun at 1200 rpm for 5 min. After blocking, attached TECs were stained with primary antibodies containing rat-anti-keratin 8 (Troma-1, DSHB, 1:20) and rabbit-anti-lamin-B1 (1:500, Abcam), or rabbit-anti-keratin 5 (Biolegend, 1:300) and mouse-anti-lamin-B1 (1:300, Santa Cruz) at room temperature (RT) for 1 hour, followed by Alexa488 goat anti-rat or anti-rabbit (for keratins) and Alexa568 goat anti-rabbit or anti-mouse (for lamin-B1) secondary antibodies. Confocal images were acquired using a laser-scanning confocal microscope (Leica SP5) with a 20X or 63X objectives. Images were processed using ImageJ.

### RNA preparation and quantitative real-time PCR

TECs were isolated and enriched as described above. Total RNA was extracted following the manufacturer’s protocol of Direct-zol™ RNA MicroPrep kit (Zymo Research R2060). Quantitative RT-PCR was performed using the iScript One Step RT-PCR kit (170-8892; Bio-Rad Laboratories) on a real-time PCR detection system (CFX96; Bio-Rad Laboratories). 50 ng of total RNA was reverse transcribed and amplified as follows: 50°C for 10 min, 95°C for 5 min, 95°C for 10 s, 60°C for 30 s, 72°C for 1 min. Steps 2–4 were repeated for 40 cycles. Each reaction was performed in triplicate, and the results of three independent experiments were used for statistical analysis. Relative mRNA expression levels were quantified using the ΔΔC (t) method (Pfaffl, 2001). Results were normalized to those for GAPDH, and primer sequences were listed as follows.

TNF-T: forward primer (F), CTGTAGCCCACGTCGTAGC;

Reverse primer (R), TTGAGATCCATGCCGTTG.

IL-1I: F, TGTAATGAAAGACGGCACACC; R, TCTTCTTTGGGTATTGCTTGG.

IL-1I: F, CCGAGT TTCATTGCCTCTTT; R, ACTGTGGGAGTGGAGTGCTT.

IL-6: F, CTCTGGGAAATCGTGGAAAT; R, CCAGTTTGGTAGCATCCATC.

P21: F, GACAAGAGGCCCAGTACTTC; R, GCTTGGAGTGATAGAAATCTGTC. P16: F, CGTACCCCG ATTCAGGTGAT; R, TTGAGCAGAAGAGCTGCTACGT. GAPDH: F, CGACTTCAACAGCAACTCCCACTCTTCC;

R, TGGGTGGTCCAGGGTTTCTTACTCCTT.

*Lmnb1*: F, AGTTTAGAGGGAGACTTGGAGG; R, TAAGGCTCTGACAGCGATTC

*Lmnb2*: F, TTAGGCCTCCAAAAGCAGG; R, CGTCATCTCCTGTTCCTTAGC

*Lmna*: F, TGAGAAGCGCACATTGGAG; R, GCGTCTGTAGCCTGTTCTC

### Western blotting analysis

Whole-cell lysates were generated using Laemmli sample buffer (Bio-rad) and diluted in SDS-PAGE sample buffer. Cell lysates were separated in 10% or 15% SDS-PAGE and then transferred onto nitrocellulose membranes. The membranes were blocked with 5% milk and probed with the following antibodies: rabbit anti-lamin-B1 (1:5000, Abcam), mouse anti-lamin-A/C (1:5000, Active Motif), rabbit anti-TNF-α (1:1000, Cell Signaling), rabbit anti-IL-1β (1:1000, Cell Signaling), rabbit anti-IL-6 (1:1000, Novus biological), rabbit anti-IL-1α (1:2000, Santa cruz), and mouse anti-β-actin (1:4000, Sigma, AC-15). Antibodies were detected with HRP-conjugated anti-mouse (1:10000) or anti-rabbit (1:10000) antibodies and West Pico Substrate (Thermo Scientific).

### Senescence-associated (SA)-β-gal assay

Senescence β-Galactosidase Staining kit (Cell Signaling, 9860S) was used to detect β-galactosidase activity at pH6.0. In brief, *in vitro* cultured TEC cells were fixed in the fixative solution for 10 min at RT. The fixed cells were washed twice with PBS and then incubated with β-galactosidase staining solution containing X-gal at 37°C overnight. After the blue color developed, bright-field cell images were taken using an Axiovert 25 microscope (Carl Zeiss) connected to a Canon camera. Total 150 cells from 3 biological replicates were counted from each experimental group for quantification.

### Tamoxifen (TAM) induced *Lmnb1* deletion

Stock solution of TAM (20 mg/ml, Sigma) was prepared by dissolving TAM powder in one volume of 100% ethanol at 55 °C for 2 min and then mixed well with 9 volumes of pre-warmed (55 °C) corn oil. To induce *Lmnb1* (exon 2) deletion, 2-mon-old mice were treated with a single intraperitoneal (IP) injection of TAM (1mg/10 g body weight/day) for five successive days and with 25 μg/ml TAM containing drinking water for the 1^st^ month (Cheng et al., 2010). Thymus phenotypes were analyzed 1 month after the last TAM injection. Peripheral T cell phenotypes were analyzed 1 month or 6 months after the last TAM injection.

### Endotoxin-induced thymic involution model

The procedure follows a detailed protocol described in (Billard et al., 2011). In brief, Escherichia coli-derived lipopolysaccharide (LPS, Sigma-Aldrich L-2880) was reconstituted at 1 mg/mL in PBS. 2-mon-old female C57BL/6 mice were injected once intraperitoneally with LPS (100 µg) or PBS to induce thymic involution. Thymuses were collected at indicated time points for phenotype analyses and lamin measurement.

### Single-cell RNA-sequencing and data processing

Immediately post-sorting, CD45^-^ DAPI^-^ EpCAM^+^ total TECs were run on the 10X Chromium and then single-cell RNA-seq libraries were generated using the Chromium Single Cell 3’ Reagent Kit (10X Genomics) by the core facility at the Embryology Department of Carnegie Institute for Science. Briefly, TEC single-cell suspension (~1,000 cells per 1 µl PBS) was mixed thoroughly with Single Cell 3’ gel beads and partitioning oil into a Single Cell 3’ Chip (10X genomics) following the recommended protocol for the Chromium Single Cell 3’ Reagent Kit (v2 Chemistry). RNA transcripts of single cells were uniquely barcoded and reverse-transcribed within the individual droplets. cDNA molecules were then pre-amplified and pooled together followed by the final library construction. Libraries were sequenced by paired-end 150-bp reads on Illumina NextSeq500. Post-processing and quality control were performed by the same genomics core facility using the 10X Cell Ranger package (V2.1.1, 10X Genomics) as described by Zheng at al. (G. X. Zheng et al., 2017). Reads were aligned to mm10 reference assembly (v1.2.0, 10X Genomics). After cell demultiplexing and read counting, all three samples were combined using the cellranger aggr pipeline (G. X. Zheng et al., 2017). Cell visualization was done using the Loupe Cell Browser (10X Genomics).

### Whole-Transcriptome Shotgun Sequencing (RNA-seq)

By 1 month after the last TAM injection, ~5000 cTECs from *Lmnb1^+/+^;K8CreERT2* and *Lmnb1^f/f^;K8CreERT2* mouse thymuses were sorted directly into Trizol by FACS sorting. Total RNA was extracted following the manufacturer’s protocol of Direct-zol™ RNA MicroPrep kit (Zymo Research). Poly-A selected mRNA was purified and sequencing libraries were built using Illumina TruSeq RNA sample prep kit V2 (Illumina). Libraries were sequenced by single end 50-bp reads on Illumina HiSeq-2000.

### Bioinformatics

For RNA-seq, low-quality reads (quality score <20) were trimmed and the trimmed reads shorter than 36 bp were then filtered out. The remaining reads were further mapped to mouse genome (UCSC, mm9) by Tophat2. The number of reads falling into each gene was then counted using the custom scripts. The differentially expressed genes were called by edgeR (Robinson et al., 2010) (fold change ≥ 2, FDR < 0.05 and p value <0.05). GO term analyses of the differentially expressed genes were performed by DAVID (Huang da et al., 2009).

## Acknowledgement

We thank Dr. Jonathan Powell, Dr. Chen-ming Fan, and the members of the Zheng lab for comments and valuable help, Dr. Youngjo Kim for generating *Lmna*^f/f^ and *Lmnb1*^-/-^ mouse alleles, Allison Pinder and Frederick Tan for RNA-Seq. This paper was supported by a senior scholar award to YZ from the Ellison Medical Foundation, NIH/NIGMS grants GM106023 (YZ) and GM110151 (YZ).

## Author contributions

Yixian Zheng and Sibiao Yue conceived the idea. Sibiao Yue performed all the experiments. Yixian Zheng and Sibiao Yue interpreted the data and wrote the paper. Xiaobin Zheng analyzed and interpreted all the RNA-seq data and performed GO term analyses of transcriptionally differentially expression of genes. The authors declare no competing financial interests.

## Supplementary Figures and Tables

**Figure S1.**
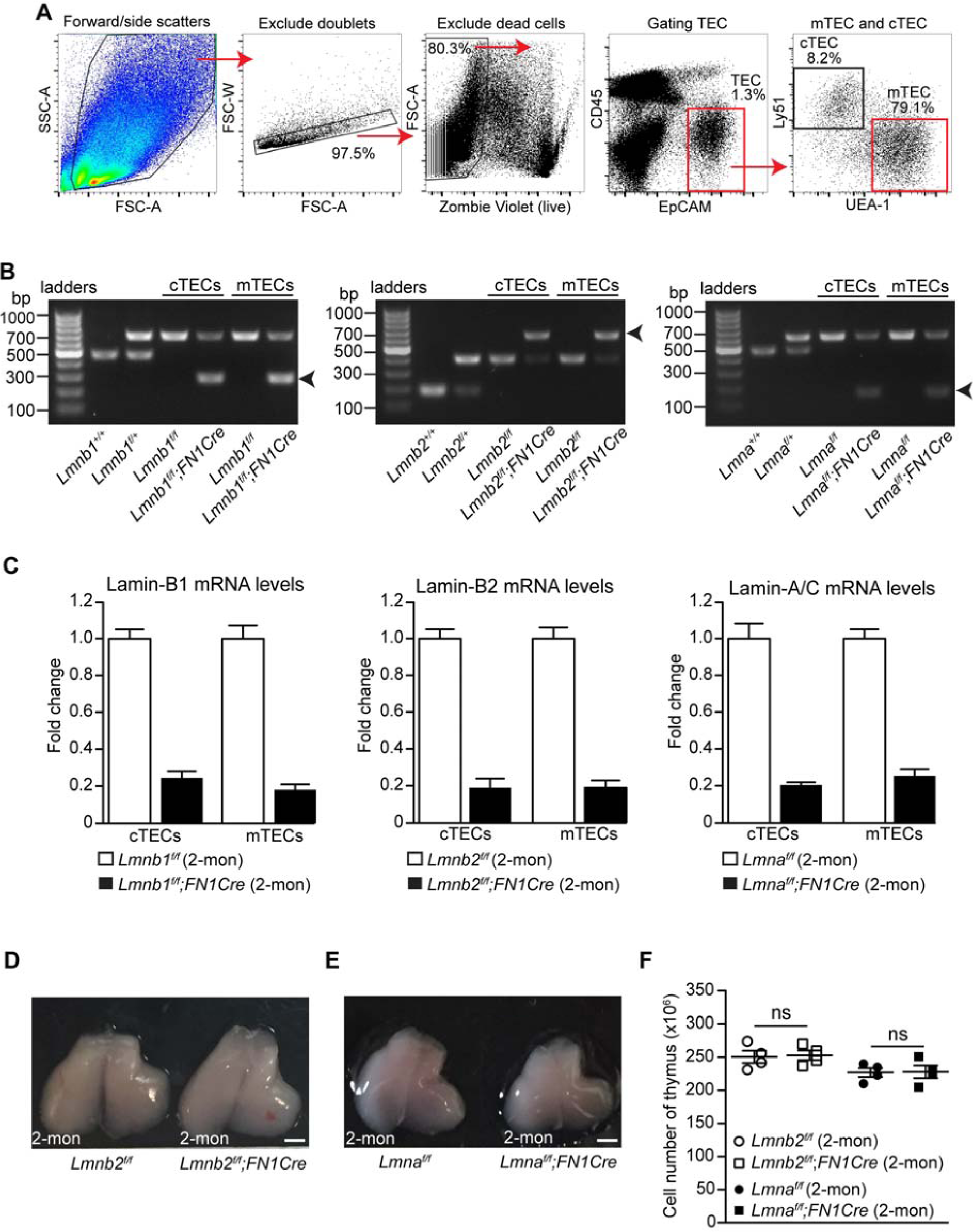
The effects of *Lmnb2* or *Lmna* deletion in TECs on thymus size and cellularity. (A) The flow cytometry gating strategy to identify cTECs and mTECs. Cells from a 2-mon-old WT thymus are shown as an example. After excluding doublet cells by forward- and side-scatter and dead cells by fixable Zombie violet dye (Biolegend), the TECs were identified first as CD45-EpCAM^+^ cells and then analyzed according to UEA-1 and Ly51 to differentiate cTECs (UEA-1-Ly51^+^) and mTECs (UEA-1^+^Ly51^-^). The percentages of the gated cells (outlined by squares) in each panel are indicated. (B) PCR genotyping of WT, *Lmnb1*^*f/f*^, *Lmnb2^f/f^, and Lmnaf/f* alleles. To test the deletion efficiency of the floxed lamin alleles, the mice were crossed with a mouse strain that express Cre driven by the *Foxn1* promoter (*FN1Cr* e) to generate TEC-specific lamin null alleles. WT, *Lmnb1*^*f*^, or *Lmnb1*^*-*^ allele generates 470bp, 670bp, or 284 bp PCR fragments, respectively. WT, *Lmnb2*^*f*^, or *Lmnb2*^*-*^ allele generates 180bp, 400bp, or 631bp PCR fragments, respectively. WT, *Lmna*^*f*^, or *Lmna*^*-*^ allele generates 470bp, 598bp, or 153bp PCR fragments, respectively. The presence of PCR products of the floxed WT lamin alleles in the *Lmnb1^f/f^;FNCre*, *Lmnb2^f/f^;FNCre*, and *Lmna^f/f^;FNCre* mice indicate the incomplete deletion of lamins. (C) RT-qPCR analyses of mRNA levels of lamin-B1, lamin-B2, and lamin-A/C in cTECs and mTECs isolated from *Lmnb1^f/f^;FN1Cre, Lmnb2f/f;FN1Cre, Lmnaf/f;FN1Cre*, and corresponding littermate controls. The lamin mRNA was normalized to GPADH mRNA level. The plots are representative results from two biological replicates. (D) Representative images of thymuses from 2-mon-old *Lmnb2*^*f/f*^ control and *Lmnb2^f/f^;FN1Cre* mice. (E) Representative images of thymuses from 2-mon-old *Lmna*^*f/f*^ control and *Lmna^f/f^;FN1Cre* mice. (F) The total thymic cell number counts from thymuses with indicated ages and genotypes. Scale bars, 1mm. Error bars, SEM from at least three independently analyzed mice. Student’s t test: *P<0.05, ** P<0.01, ***P<0.001. ns: not significant.

**Figure S2.**
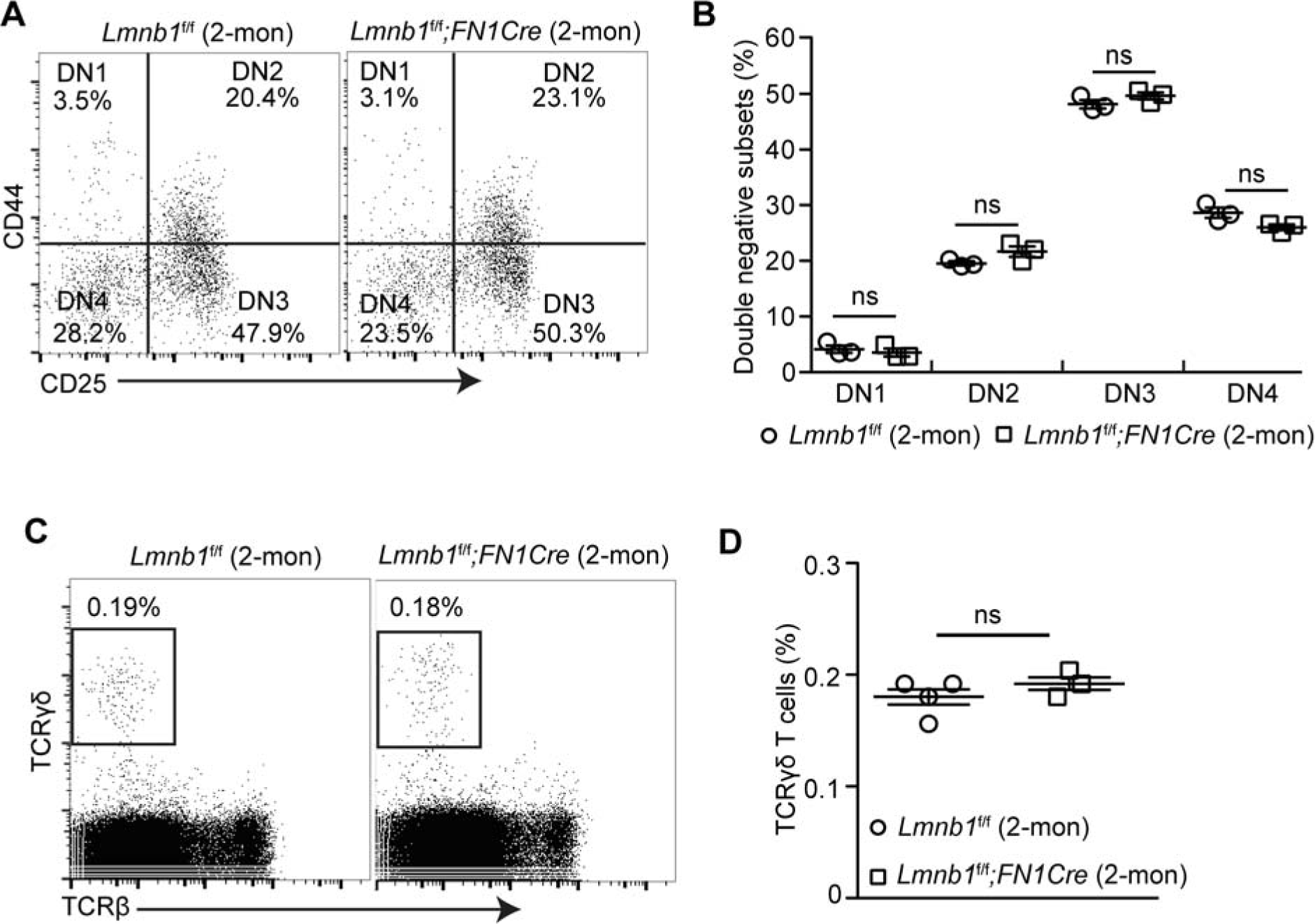
Effects of embryonic *Lmnb1* deletion in TECs on double negative and γδT cell development. (A) Flow cytometry analyses of CD44 and CD25 expression on the CD4^-^CD8^-^ double-negative (DN) thymocyte population in 2-mon-old *Lmnb1*^*f/f*^ control and *Lmnb1^f/f^;FN1Cre* mouse thymuses. The DN cells are subdivided into DN1 (CD44^+^CD25^-^), DN2 (CD44^+^CD25^+^), DN3 (CD44^-^CD25^+^), and DN4 (CD44^-^CD25^-^) based on differential expression of CD44 and CD25. (B) Quantification of the frequency of DN1-4 subsets in the 2-mon-old *Lmnb1*^*f/f*^ control (n=3) and *Lmnb1^f/f^;FN1Cre* (n=3) mouse thymuses. (C) Flow cytometry analyses of TCR and TCR expression on thymocytes in 2-mon-old *Lmnb1*^*f/f*^ control and *Lmnb1^f/f^;FN1Cre* mouse thymuses. (D) Quantification of the frequency of ((T cells in the 2-mon-old control *Lmnb1*^*f/f*^ (n=4) and *Lmnb1^f/f^;FN1Cre* thymus (n=3). Error bars, SEM from at least 3 mice for each genotype. Student’s t test: *P<0.05, **P< 0.01, ***P<0.001, ns: not significant.

**Figure S3.**
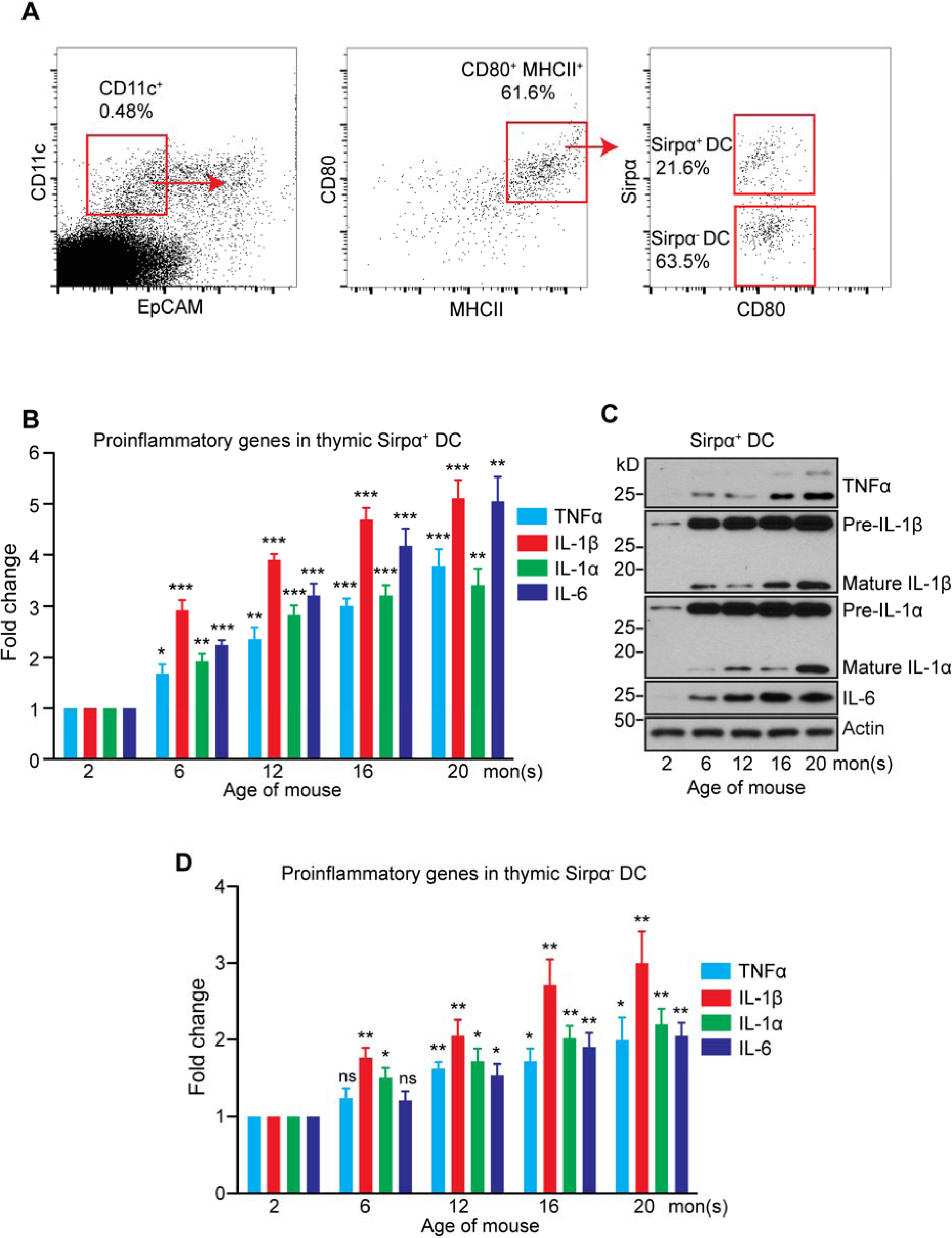
Age-associated increase in proinflammatory cytokines in Sirpα^+^ dendritic cells (DC) (A) The flow cytometry gating strategy for sorting Sirpα^+^ and Sirpα^-^ DCs from a 2-mon-old WT thymus. DCs were identified based on the expression of CD11c, MHCII, and CD80. (B) RTqPCR analyses of TNF-α, IL-1β, IL-1α, and IL-6 in Sirpα^+^ DCs isolated from WT thymuses at the indicated ages. The increased expression was plotted relative to that of the Sirpα^+^ DCs isolated from 2-mon-old thymus, which was set to 1. (C) The representative Western blots of TNF-α, IL-1β, IL-1α, and IL-6 in Sirpα^+^ DCs isolated from WT thymuses at the indicated ages. β-actin, loading control. (D) RT-qPCR analyses of TNF-α, IL-1β, IL-1α, and IL-6 in Sirpα^-^ DCs isolated from WT thymuses at the indicated ages. The increased expression was plotted relative to that of the Sirpα^-^ DCs isolated from 2-mon-old thymus, which was set to 1. Error bars, SEM from three independent experiments. Student’s t test: *P<0.05, **P<0.01, ***P<0.001, ns: not significant.

**Figure S4.**
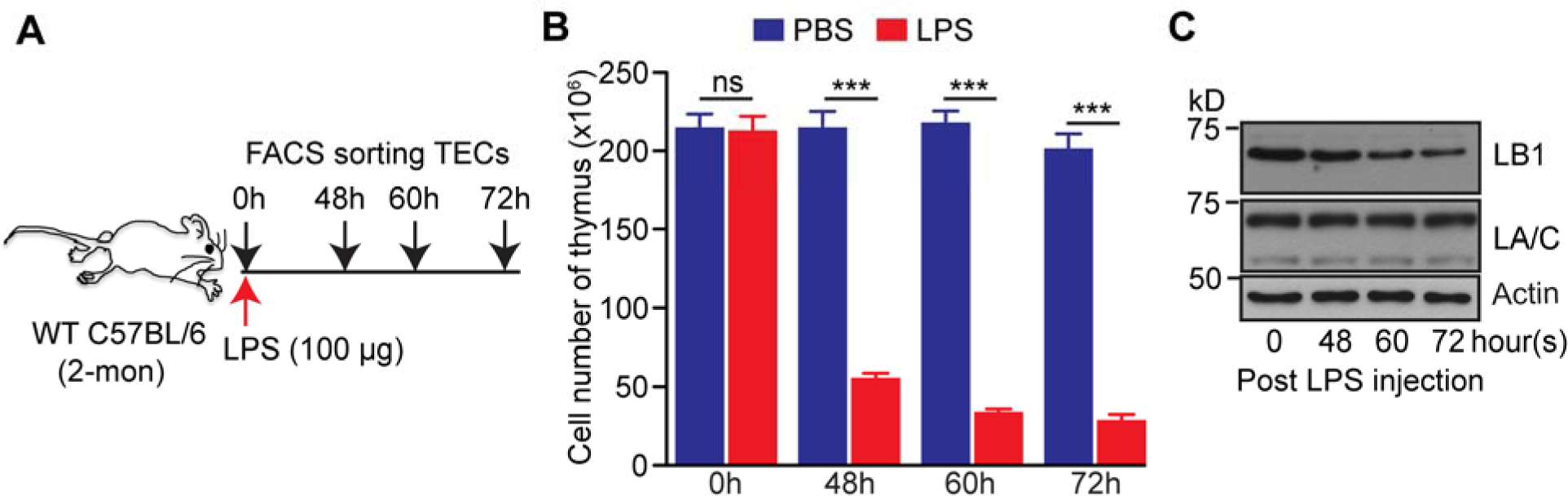
Effects of acute thymic inflammation induced by lipopolysaccharide (LPS) on thymic atrophy and TEC lamin-B1 levels. (A) Schematic depiction of LPS-induced thymic involution model. Intraperitoneal injection (IP) of one-dose of LPS (100 tg) was used to induce acute thymic involution. (B) Total cell numbers of thymuses at the indicated time points post LPS challenge. (C) Western blotting analyses of lamin-B1 and lamin-A/C in TECs collected by FACS sorting from 3 pooled thymuses at the indicated time of LPS injection. β-actin is used as a loading control. Error bars, SEM from at least three independent experiments. Student’s t test: *P<0.05, **P<0.01, ***P<0.001, ns: not significant.

**Figure S5.**
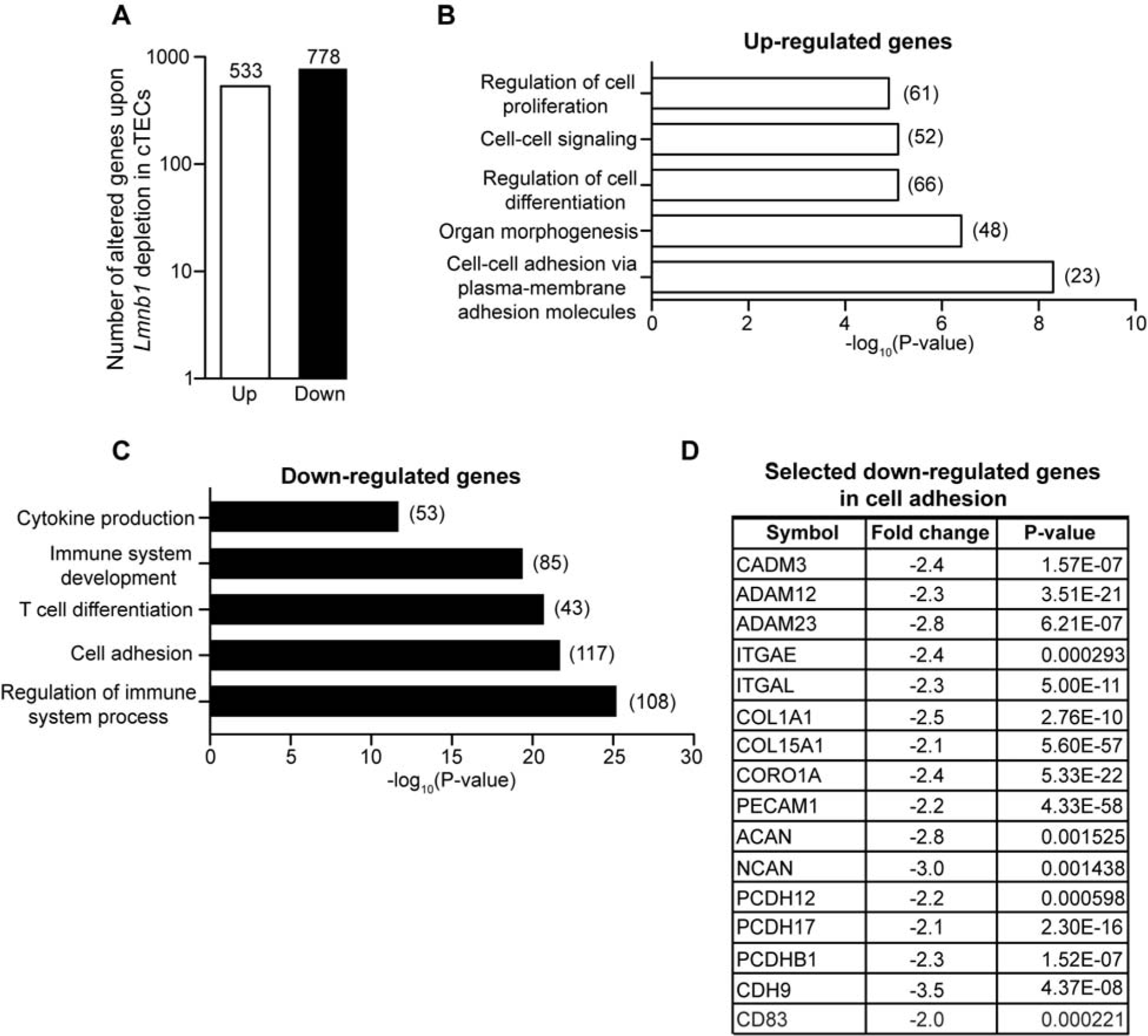
Effects of postnatal lamin-B1 deficiency in cTECs on gene expression. (A) A plot of the number of up- (white) and down-regulated (black) genes (fold change ≥2) upon postnatal *Lmnb1* deletion in cTECs (1 month after the TAM injections, see Figure 6A). (B-C) DAVID GO term analyses of up-regulated (B) or down-regulated (C) genes in cTECs upon *Lmnb1* deletion. The number of altered genes in each Go term category is indicated in parentheses. (D) Selected down-regulated genes in the cell adhesion GO term category. See also Table S1 and S2.

**Figure S6.**
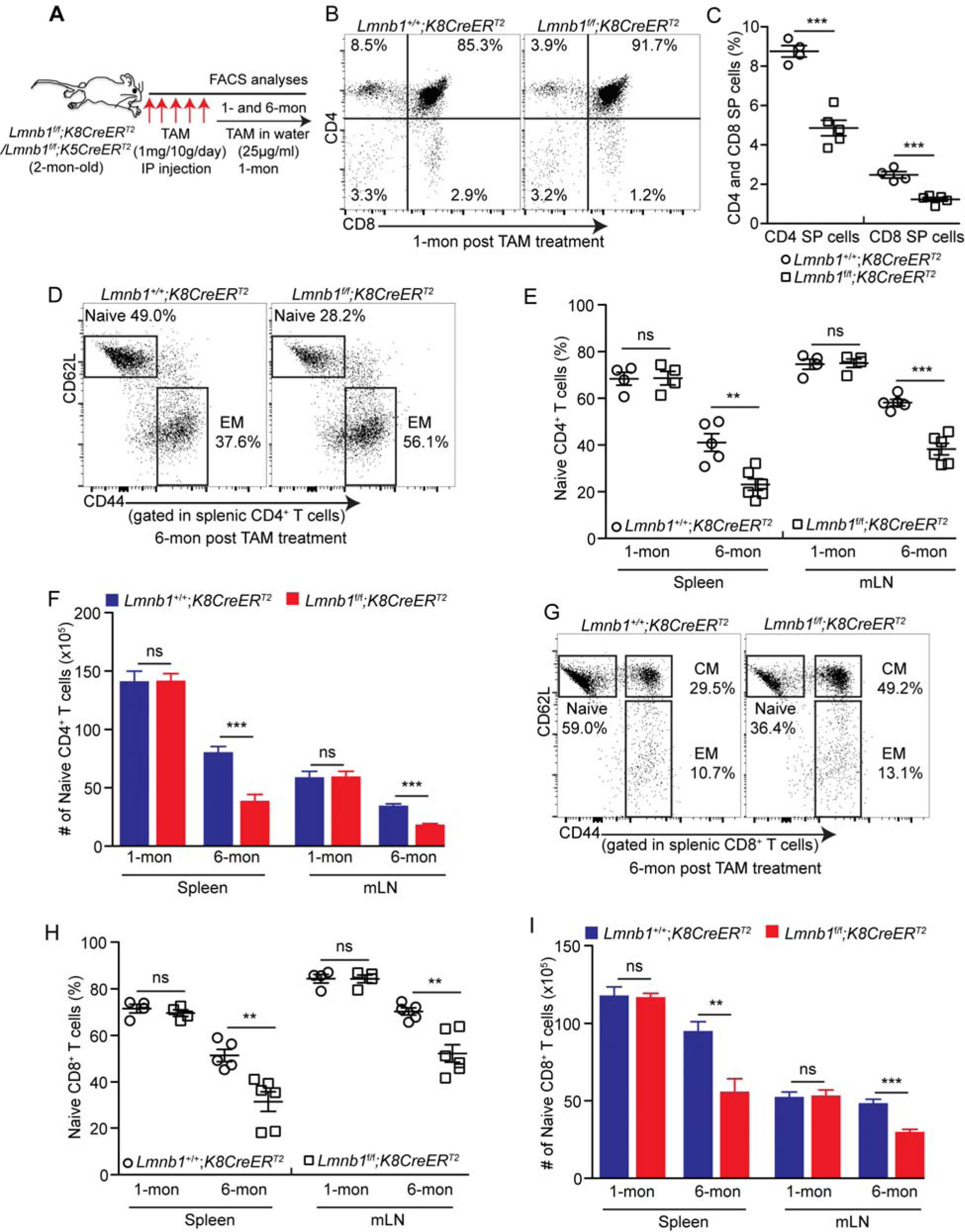
Effects of postnatal lamin-B1 deficiency in TECs on thymic function. (A) The TAM-mediated *Lmnb1* deletion scheme. Keratin 8 (*K8*)- or keratin 5 (*K5*)-driven *CreER*^*T2*^ (see Figure S6) was used to delete *Lmnb1* in cTECs or mTECs, respectively, by injecting TAM followed by feeding TAM-containing drinking water for one month followed by analyses at one or six months after the last TAM injection. (B) Representative flow cytometry profiles of CD4^+^ SP or CD8^+^ SP thymocytes from *Lmnb1^+/+^;K8CreERT2* or *Lmnb1^f/f^;K8CreERT2* thymuses 1 month after the last TAM injection. (C) Quantification of the frequency of CD4^+^ and CD8^+^ SP thymocytes among the gated live cells in *Lmnb1*^*+/+*^;*K8CreER*^*T2*^ (n=4) or *Lmnb1^f/f^;K8CreERT2* (n=5) thymuses. (D) Representative flow cytometry analyses showing a reduction of naïve CD4^+^ T cells and an increase of effector memory (EM) CD4^+^ T cells in the spleen of *Lmnb1^f/f^;K8CreERT2* mice compared to the control *Lmnb1*^*+/+*^; *K8CreER*^*T2*^ mice 6-month after the last TAM injections. The splenocytes were stained with CD3, CD4, CD62L and CD44 to identify the naïve (CD3^+^CD4^+^CD62L^+^CD44^low/-^) and EM (CD3^+^CD4^+^CD62L^-^CD44^hi^) CD4^+^ T cells. (E-F) Quantification of the frequency (E) and cell number (F) of the naïve CD4^+^ T cells in the spleen and mesenteric lymph node (mLN) in *Lmnb1^+/+^;K8CreERT2* and *Lmnb1^f/f^;K8CreERT2* mice 1 and 6 month after TAM injections. (G) Representative flow cytometry analyses showing a reduction of naïve CD8^+^ T cells and an increase of effector memory (EM) and central memory (CM) CD8^+^ T cells in the spleen of *Lmnb1^f/f^;K8CreERT2* mice compared to the control *Lmnb1*^*+/+*^; *K8CreER*^*T2*^ mice 6-month after the TAM injections. The splenocytes were stained with CD3, CD8, CD62L and CD44 to identify the naïve (CD3^+^CD8^+^CD62L^+^CD44^low/-^), EM (CD3^+^CD8^+^CD62L^-^CD44^hi^) and CM (CD3^+^CD8^+^CD62L^hi^CD44^hi^) CD8^+^ T cells. (H-I) Quantification of the frequency (H) and cell number (I) of the naïve CD8^+^ T cells in the spleen and mLN in *Lmnb1^+/+^;K8CreERT2* and *Lmnb1^f/f^;K8CreERT2* mice 1 and 6 months after the TAM injections. Each circle or square represents one *Lmnb1* control or mutant mouse. Error bars, SEM from at least three mice. Student’s t test: *P< 0.05, **P < 0.01, ***P< 0.001, ns: not significant.

**Table S1. List of genes that are changed by at least 2 Fold in the TAM-mediated *Lmnb1-* depleted cTECs compared to controls. Related to Figure S5**

Gene symbol, gene location, fold change (logFC, log2 of the fold change), p values (by hypergeometric test), and FDR (False Discover Rate) are indicated.

**Table S2. List of DAVID GO term category of up-regulated and down-regulated genes in the TAM-mediated *Lmnb1*-depleted cTECs compared to controls. Related to Figure S5**

Category term, count, fold enrichment, p values, and FDR are indicated.

**Table S3. List 1, genes that are differentially enriched in different TEC clusters identified by single-cell RNA-sequencing. List 2, percentages of each cluster present in WT young, WT old, and lamin-B1 depleted TECs. Related to Figure 6.**

Gene ID, gene name, Mean Unique Molecular Identifier (UMI) counts, fold change (logFC, log2 of the fold change), adjusted p values are indicated.

## Reference

Acosta, J. C., O’Loghlen, A., Banito, A., Guijarro, M. V., Augert, A., Raguz, S., … Gil, J. (2008). Chemokine signaling via the CXCR2 receptor reinforces senescence. Cell, 133(6), 1006–1018. doi:10.1016/j.cell.2008.03.038

Anderson, G., & Takahama, Y. (2012). Thymic epithelial cells: working class heroes for T cell development and repertoire selection. Trends Immunol, 33(6), 256–263. doi:10.1016/j.it.2012.03.005

Anderson, M. S., Venanzi, E. S., Klein, L., Chen, Z., Berzins, S. P., Turley, S. J., … Mathis, D. (2002). Projection of an immunological self shadow within the thymus by the aire protein. Science, 298(5597), 1395–1401. doi:10.1126/science.1075958

Aw, D., & Palmer, D. B. (2012). It’s not all equal: a multiphasic theory of thymic involution. Biogerontology, 13(1), 77–81. doi:10.1007/s10522–011–9349–0

Aw, D., Silva, A. B., Maddick, M., von Zglinicki, T., & Palmer, D. B. (2008). Architectural changes in the thymus of aging mice. Aging Cell, 7(2), 158–167. doi:10.1111/j.1474–9726.2007.00365.x

Billard, M. J., Gruver, A. L., & Sempowski, G. D. (2011). Acute endotoxin-induced thymic atrophy is characterized by intrathymic inflammatory and wound healing responses. PLoS One, 6(3), e17940. doi:10.1371/journal.pone.0017940

Boehm, T., Nehls, M., & Kyewski, B. (1995). Transcription factors that control development of the thymic microenvironment. Immunol Today, 16(12), 555–556. doi:10.1016/0167–5699(95)80074–3

Campisi, J. (2013). Aging, cellular senescence, and cancer. Annu Rev Physiol, 75, 685–705. doi:10.1146/annurev-physiol-030212–183653

Chen, H., Zheng, X., & Zheng, Y. (2014). Age-associated loss of lamin-B leads to systemic inflammation and gut hyperplasia. Cell, 159(4), 829–843. doi:10.1016/j.cell.2014.10.028

Chen, L., Xiao, S., & Manley, N. R. (2009). Foxn1 is required to maintain the postnatal thymic microenvironment in a dosage-sensitive manner. Blood, 113(3), 567–574. doi:10.1182/blood-2008–05–156265

Cheng, L., Guo, J., Sun, L., Fu, J., Barnes, P. F., Metzger, D., … Su, D. M. (2010). Postnatal tissue-specific disruption of transcription factor FoxN1 triggers acute thymic atrophy. J Biol Chem, 285(8), 5836–5847. doi:10.1074/jbc.M109.072124

Chien, Y. H., Meyer, C., & Bonneville, M. (2014). gammadelta T cells: first line of defense and beyond. Annu Rev Immunol, 32, 121–155. doi:10.1146/annurev-immunol-032713–120216

Chinn, I. K., Blackburn, C. C., Manley, N. R., & Sempowski, G. D. (2012). Changes in primary lymphoid organs with aging. Semin Immunol, 24(5), 309–320. doi:10.1016/j.smim.2012.04.005

Coffinier, C., Jung, H. J., Nobumori, C., Chang, S., Tu, Y., Barnes, R. H., 2nd, … Young, S. G. (2011). Deficiencies in lamin B1 and lamin B2 cause neurodevelopmental defects and distinct nuclear shape abnormalities in neurons. Mol Biol Cell, 22(23), 4683–4693. doi:10.1091/mbc.E11–06–0504

Coppe, J. P., Patil, C. K., Rodier, F., Sun, Y., Munoz, D. P., Goldstein, J., … Campisi, J. (2008). Senescence-associated secretory phenotypes reveal cell-nonautonomous functions of oncogenic RAS and the p53 tumor suppressor. PLoS Biol, 6(12), 2853–2868. doi:10.1371/journal.pbio.0060301

Corbeaux, T., Hess, I., Swann, J. B., Kanzler, B., Haas-Assenbaum, A., & Boehm, T. (2010). Thymopoiesis in mice depends on a Foxn1-positive thymic epithelial cell lineage. Proc Natl Acad Sci U S A, 107(38), 16613–16618. doi:10.1073/pnas.1004623107

Depreter, M. G., Blair, N. F., Gaskell, T. L., Nowell, C. S., Davern, K., Pagliocca, A., … Blackburn, C. C. (2008). Identification of Plet-1 as a specific marker of early thymic epithelial progenitor cells. Proc Natl Acad Sci U S A, 105(3), 961–966. doi:10.1073/pnas.0711170105

Dreesen, O., Chojnowski, A., Ong, P. F., Zhao, T. Y., Common, J. E., Lunny, D., … Colman, A. (2013). Lamin B1 fluctuations have differential effects on cellular proliferation and senescence. J Cell Biol, 200(5), 605–617. doi:10.1083/jcb.201206121

Dumont, P., Balbeur, L., Remacle, J., & Toussaint, O. (2000). Appearance of biomarkers of in vitro ageing after successive stimulation of WI-38 fibroblasts with IL-1alpha and TNF-alpha: senescence associated beta-galactosidase activity and morphotype transition. J Anat, 197 Pt 4, 529–537.

Franceschi, C., Bonafe, M., Valensin, S., Olivieri, F., De Luca, M., Ottaviani, E., & De Benedictis, G. (2000). Inflamm-aging. An evolutionary perspective on immunosenescence. Ann N Y Acad Sci, 908, 244–254.

Frost, B., Bardai, F. H., & Feany, M. B. (2016). Lamin Dysfunction Mediates Neurodegeneration in Tauopathies. Curr Biol, 26(1), 129–136. doi:10.1016/j.cub.2015.11.039

Gordon, J., Xiao, S., Hughes, B., 3rd, Su, D. M., Navarre, S. P., Condie, B. G., & Manley, N. R. (2007). Specific expression of lacZ and cre recombinase in fetal thymic epithelial cells by multiplex gene targeting at the Foxn1 locus. BMC Dev Biol, 7, 69. doi:10.1186/1471–213X-7–69

Hennet, T., Hagen, F. K., Tabak, L. A., & Marth, J. D. (1995). T-cell-specific deletion of a polypeptide N-acetylgalactosaminyl-transferase gene by site-directed recombination. Proc Natl Acad Sci U S A, 92(26), 12070–12074.

Huang da, W., Sherman, B. T., & Lempicki, R. A. (2009). Bioinformatics enrichment tools: paths toward the comprehensive functional analysis of large gene lists. Nucleic Acids Res, 37(1), 1–13. doi:10.1093/nar/gkn923

Jain, R., & Gray, D. H. (2014). Isolation of thymic epithelial cells and analysis by flow cytometry. Curr Protoc Immunol, 107, 3 26 21–15. doi:10.1002/0471142735.im0326s107

Kernfeld, E. M., Genga, R. M. J., Neherin, K., Magaletta, M. E., Xu, P., & Maehr, R. (2018). A Single-Cell Transcriptomic Atlas of Thymus Organogenesis Resolves Cell Types and Developmental Maturation. Immunity. doi:10.1016/j.immuni.2018.04.015

Ki, S., Park, D., Selden, H. J., Seita, J., Chung, H., Kim, J., … Ehrlich, L. I. (2014). Global transcriptional profiling reveals distinct functions of thymic stromal subsets and age-related changes during thymic involution. Cell Rep, 9(1), 402–415. doi:10.1016/j.celrep.2014.08.070

Kim, Y., Sharov, A. A., McDole, K., Cheng, M., Hao, H., Fan, C. M., … Zheng, Y. (2011). Mouse B-type lamins are required for proper organogenesis but not by embryonic stem cells. Science, 334(6063), 1706–1710. doi:10.1126/science.1211222

Kim, Y., & Zheng, Y. (2013). Generation and characterization of a conditional deletion allele for Lmna in mice. Biochem Biophys Res Commun, 440(1), 8–13. doi:10.1016/j.bbrc.2013.08.082

Klug, D. B., Carter, C., Crouch, E., Roop, D., Conti, C. J., & Richie, E. R. (1998). Interdependence of cortical thymic epithelial cell differentiation and T-lineage commitment. Proc Natl Acad Sci U S A, 95(20), 11822–11827.

Kuilman, T., Michaloglou, C., Vredeveld, L. C., Douma, S., van Doorn, R., Desmet, C. J., … Peeper, D. S. (2008). Oncogene-induced senescence relayed by an interleukin-dependent inflammatory network. Cell, 133(6), 1019–1031. doi:10.1016/j.cell.2008.03.039

Miragaia, R. J., Zhang, X., Gomes, T., Svensson, V., Ilicic, T., Henriksson, J., … Lonnberg, T. (2018). Single-cell RNA-sequencing resolves self-antigen expression during mTEC development. Sci Rep, 8(1), 685. doi:10.1038/s41598–017–19100–4

O’Neill, K. E., Bredenkamp, N., Tischner, C., Vaidya, H. J., Stenhouse, F. H., Peddie, C. D., … Blackburn, C. C. (2016). Foxn1 Is Dynamically Regulated in Thymic Epithelial Cells during Embryogenesis and at the Onset of Thymic Involution. PLoS One, 11(3), e0151666. doi:10.1371/journal.pone.0151666

Ren, J. L., Pan, J. S., Lu, Y. P., Sun, P., & Han, J. (2009). Inflammatory signaling and cellular senescence. Cell Signal, 21(3), 378–383. doi:10.1016/j.cellsig.2008.10.011

Robinson, M. D., McCarthy, D. J., & Smyth, G. K. (2010). edgeR: a Bioconductor package for differential expression analysis of digital gene expression data. Bioinformatics, 26(1), 139–140. doi:10.1093/bioinformatics/btp616

Rode, I., Martins, V. C., Kublbeck, G., Maltry, N., Tessmer, C., & Rodewald, H. R. (2015). Foxn1 Protein Expression in the Developing, Aging, and Regenerating Thymus. J Immunol, 195(12), 5678–5687. doi:10.4049/jimmunol.1502010

Rodier, F., & Campisi, J. (2011). Four faces of cellular senescence. J Cell Biol, 192(4), 547–556. doi:10.1083/jcb.201009094

Shanley, D. P., Aw, D., Manley, N. R., & Palmer, D. B. (2009). An evolutionary perspective on the mechanisms of immunosenescence. Trends Immunol, 30(7), 374–381. doi:10.1016/j.it.2009.05.001

St-Pierre, C., Brochu, S., Vanegas, J. R., Dumont-Lagace, M., Lemieux, S., & Perreault, C. (2013). Transcriptome sequencing of neonatal thymic epithelial cells. Sci Rep, 3, 1860. doi:10.1038/srep01860

Tran, J. R., Chen, H., Zheng, X., & Zheng, Y. (2016). Lamin in inflammation and aging. Curr Opin Cell Biol, 40, 124–130. doi:10.1016/j.ceb.2016.03.004

Ulyanchenko, S., O’Neill, K. E., Medley, T., Farley, A. M., Vaidya, H. J., Cook, A. M., … Blackburn, C. C. (2016). Identification of a Bipotent Epithelial Progenitor Population in the Adult Thymus. Cell Rep, 14(12), 2819–2832. doi:10.1016/j.celrep.2016.02.080

Youm, Y. H., Kanneganti, T. D., Vandanmagsar, B., Zhu, X., Ravussin, A., Adijiang, A., … Dixit, V. D. (2012). The Nlrp3 inflammasome promotes age-related thymic demise and immunosenescence. Cell Rep, 1(1), 56–68. doi:10.1016/j.celrep.2011.11.005

Zheng, G. X., Terry, J. M., Belgrader, P., Ryvkin, P., Bent, Z. W., Wilson, R., … Bielas, J. H. (2017). Massively parallel digital transcriptional profiling of single cells. Nat Commun, 8, 14049. doi:10.1038/ncomms14049

Zheng, X., Hu, J., Yue, S., Kristiani, L., Kim, M., Sauria, M., … Zheng, Y. (2018). Lamins Organize the Global Three-Dimensional Genome from the Nuclear Periphery. Mol Cell. doi:10.1016/j.molcel.2018.05.017

Zook, E. C., Krishack, P. A., Zhang, S., Zeleznik-Le, N. J., Firulli, A. B., Witte, P. L., & Le, P. T. (2011). Overexpression of Foxn1 attenuates age-associated thymic involution and prevents the expansion of peripheral CD4 memory T cells. Blood, 118(22), 5723–5731. doi:10.1182/blood-2011–03–342097

